# A novel type of monocytic leukemia stem cell revealed by the clinical use of venetoclax-based therapy

**DOI:** 10.1101/2022.12.04.519036

**Authors:** Shanshan Pei, Austin E Gillen, Ian T Shelton, Brett M Stevens, Maura Gasparetto, Krysta Engel, Sarah Staggs, Yanan Wang, William Showers, Anagha Inguva, Maria L Amaya, Mohammad Minhajuddin, Amanda Winters, Sweta B Patel, Hunter Tolison, Anna Krug, Tracy N Young, Jeffrey Schowinsky, Christine McMahon, Clayton A Smith, Daniel A Pollyea, Craig T Jordan

**Affiliations:** Bone Marrow Transplantation Center, the First Affiliated Hospital, School of Medicine, Zhejiang University, Hangzhou, China; Liangzhu Laboratory, Zhejiang University Medical Center, Hangzhou, China; Division of Hematology, University of Colorado School of Medicine, Aurora, Colorado, USA; Institute of Hematology, Zhejiang University, Hangzhou, China; Zhejiang Province Engineering Laboratory for Stem Cell and Immunity Therapy, Hangzhou, China; Center for Cancer and Blood Disorders, Department of Pediatrics, University of Colorado, Aurora, Colorado, USA

**Keywords:** Acute myeloid leukemia, Leukemia stem cells, Monocytic AML, Relapse, Venetoclax, Cladribine, Heterogeneity

## Abstract

The BCL-2 inhibitor venetoclax has recently emerged as an important component of acute myeloid leukemia (AML) therapy. Notably, use of this agent has revealed a previously unrecognized form of pathogenesis characterized by monocytic disease progression. We demonstrate that this form of disease arises from a fundamentally different type of leukemia stem cell (LSC), which we designate as monocytic LSC (m-LSC), that is developmentally and clinically distinct from the more well-described primitive LSC (p-LSC). The m-LSC is distinguished by a unique immunophenotype (CD34-, CD4+, CD11b-, CD14-, CD36-), unique transcriptional state, reliance on purine/pyrimidine metabolism, and selective sensitivity to cladribine. Critically, in some instances m-LSC and p-LSC subtypes can co-reside in the same AML patient and simultaneously contribute to overall tumor complexity. Thus, our findings demonstrate that LSC heterogeneity has direct clinical significance and highlights the need to distinguish and target m-LSCs as a means to improve clinical outcomes with venetoclax-based regimens.

**Statement of Significance:** These studies identify and characterize a new type of human acute myeloid leukemia stem cell (LSC) that is responsible for monocytic disease progression in acute myeloid leukemia (AML) patients treated with venetoclax-based regimens. Our studies describe the phenotype, molecular properties, and drug sensitivities of this unique LSC subclass.

## Introduction

Numerous studies have described the properties of malignant stem cells that drive the pathogenesis of myeloid leukemias (1). Analyses of primary human tissue specimens, as well as many different mouse models have consistently shown that leukemia stem cells (LSCs) are biologically distinct from bulk tumor populations, and frequently demonstrate drug sensitivity/resistance profiles that differ from the majority of leukemic cell types (2). Analogous to normal hematopoietic stem cells, conventional LSCs are also thought to be mostly quiescent, and capable of giving rise to progeny that comprise the overall tumor population. As such, LSCs represent a critical target in the development of novel therapies. Previous attempts to target the LSC population have been made, including focusing on specific cell surface antigens, metabolic interventions, epigenetic strategies, mutation-targeted approaches, immune-based therapies, and more (2). While multiple strategies derive from robust experimental evidence, as yet improvement of clinical outcomes due to direct eradication of LSCs has remained limited.

We posit that a major challenge in targeting LSCs lies in the inherent heterogeneity of malignant stem cells (3). As reported by multiple previous studies, LSC populations derived from human AML patients demonstrate significant intra- and inter-patient heterogeneity in developmental stages and immunophenotypes (4-9). Furthermore, single cell studies have begun to describe underlying genetic diversity that may contribute to LSC heterogeneity (10, 11). Thus, to improve clinical outcomes from LSC-targeting therapies, it is likely critical to understand the properties of heterogeneous LSC subtypes and employ therapeutic strategies that either target common vulnerabilities or are designed to simultaneously eradicate differing LSC types.

To better elucidate the nature of LSC heterogeneity we recently described analysis of patients who relapse following treatment with the recently FDA-approved regimen containing the BCL-2 inhibitor venetoclax in combination with azacitidine (12). These studies demonstrated two key findings. First, we observed that relapsed disease can in some cases manifest with a more differentiated monocytic phenotype, specifically those with AML categorized by the French American British (FAB) system as M5 but not M4 (12, 13). This finding is quite distinct from patients treated with conventional chemotherapy who almost invariably relapse with disease that shows a more primitive phenotype (12, 14). Second, these studies suggest that stem cells driving monocytic relapse may be biologically distinct from the LSCs that create more primitive de novo disease. Together, these findings indicate the existence of heterogeneous underlying LSC populations that mediate differing therapeutic outcomes of conventional chemotherapy and venetoclax-based therapies. In the present study, we performed an in-depth analysis of human AML patient specimens as a means to identify and characterize LSC heterogeneity that may drive the evolution of disease in response to venetoclax-based regimens.

## Results

### Characterization of developmentally heterogeneous LSCs

As a resource for the studies described herein, we first analyzed a cohort of 20 primary human patient specimens that were selected to represent the spectrum of phenotypes commonly encountered in AML. As illustrated in **Fig. 1A**, we defined three broad classes of specimens: 1) predominantly primitive (Prim), 2) mixed monocytic-primitive (MMP), and 3) predominantly monocytic (Mono). As indicated by the nomenclature, the Prim specimens display a leukemia blast-like profile (CD45-medium, SSC-medium), high expression of stem/progenitor associated markers (CD34 and CD117), and low expression of monocytic antigens (CD11b, CD64, CD14, CD36, and LILRB4), representing the left end of the myeloid developmental spectrum (**Fig. 1A**). In contrast, on the right end of the spectrum, Mono specimens display a monocyte-like profile (CD45-bright, SSC-high), down-regulate CD34 and CD117, and up-regulate monocytic markers to varying degrees. Finally, the MMP group contains a mixture of monocytic and primitive cells, occupying the middle range of the developmental spectrum (please see **Supplementary Fig. 1A-C** and **Supplementary Table S1** for further details).

**Fig.1.**
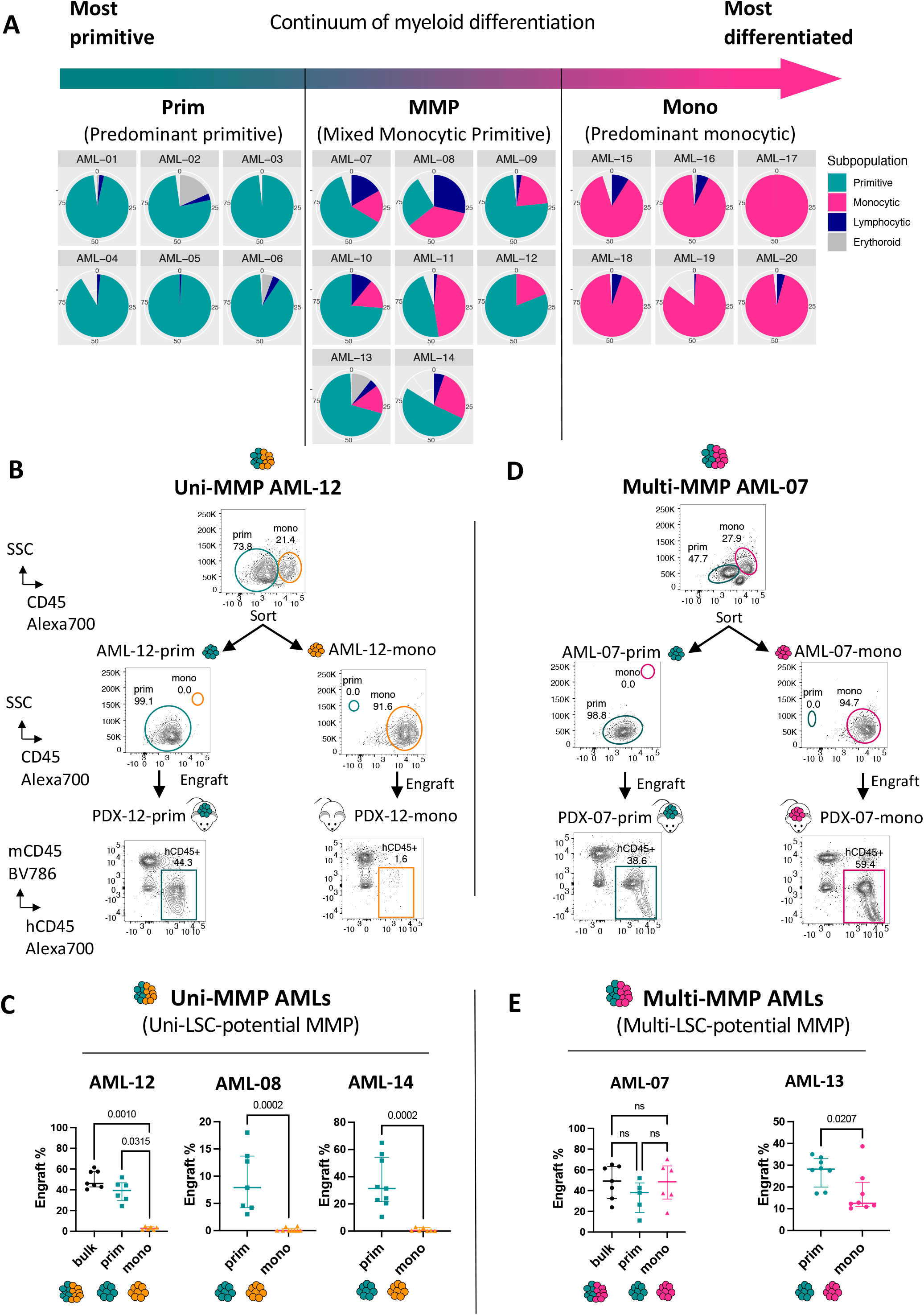
Characterization of developmentally heterogeneous LSCs. **A**, Pie charts showing relative proportion of cells resting at primitive (teal), monocytic (pink), lymphocytic (dark blue), and erythroid (gray) stages for each primary AML (Supplementary Fig. 1 and Supplementary Table S1). **B, D**, Sorting strategy and engraftment of prim and mono subpopulations from representative Uni-MMP AML-12 and Multi-MMP AML-07. The CD45/SSC flow plots in the top row depict leukemia disease before sort; The CD45/SSC flow plots in the middle demonstrate cells post sort; The human-CD45/mouse-CD45 (hCD45/mCD45) flow plots in the bottom row show engraftment levels in bone marrow of representative recipient mice. Percentage of hCD45+/mCD45- cells were used to quantify engraftment levels. **C, E**, Summary of engraft% data for Uni-MMP AML-12, AML-08, and AML-14 and Multi-MMP AML-07, and AML-13. Bulk stands for unsorted bulk tumor; prim stands for primitive subpopulation; mono stands for monocytic subpopulation. Each dot represents a mouse. AML-12 (bulk, n=7; prim, n=6; mono, n=6). AML-08 (prim, n=7; mono, n=9). AML-14 (prim, n=9; mono, n=7). AML-07 (bulk, n=7; prim, n=5; mono, n=6). AML-13 (prim, n=8; mono, n=8). Median+/- interquartile range. Two-tailed Mann-Whitney tests were used for comparing two groups, Kruskal-Wallis tests were used when comparing more than two groups. ns, not significant.

We hypothesized that heterogeneity within the LSC compartment may contribute to the developmental heterogeneity illustrated in **Fig. 1A**. To begin to characterize potential LSC heterogeneity, we performed flow cytometric cell sorting to isolate primitive (prim) and monocytic (mono) subpopulations from several MMP AMLs. Each subpopulation was independently transplanted into immunodeficient NSG-S mice and evaluated for engraftment of leukemic disease by measuring the percentage of human CD45+ cells in the bone marrow of each experimental animal. These studies revealed two subgroups of MMP AML patients. One we term “Uni-MMP” where LSC activity was exclusively detected in the prim but not in the mono subpopulation (**Fig. 1B-C**). In contrast, the second type of MMP AML we term “Multi-MMP” where readily detectable LSC activity was evident in both prim and mono subpopulations (**Fig. 1D-E**). These findings corroborate and extend several previous studies demonstrating that phenotypically distinct subpopulations of LSCs can simultaneously exist in certain AML patients(6-9).

To further characterize the leukemogenic properties of LSCs in Multi-MMP patients, the nature of disease arising in transplanted NSG-S mice was evaluated. As shown in **Fig. 2A** and **Supplementary Fig. 2A-B**, prim vs mono engrafted cells from two representative specimens (AML-07 and AML-13) consistently differed in their developmental spectrum. For both AMLs, the prim subpopulation was able to recapitulate the full developmental spectrum of disease, with both prim and mono cells evident in each transplanted mouse. In contrast, the mono subpopulation only gave rise to monocytic disease, with no evidence of more primitive cells. Importantly, such distinct differences in developmental spectrum of engrafted disease also translated into clear differences in therapeutic sensitivity. When prim vs mono engrafted groups were treated with a regimen of venetoclax plus azacitidine (VEN+AZA) *in vivo*, the mono derived disease was significantly more resistant to VEN+AZA than the prim cells (**Fig. 2B-D**). Thus, the LSCs that initiate and drive disease in the monocytic subpopulation demonstrate a more restricted developmental hierarchy that resides toward the mature end of the myeloid developmental spectrum and distinct resistance to VEN+AZA therapy. We designate this subclass of LSCs as mono-LSCs (m-LSCs) in contrast to the more conventional prim-LSCs (p-LSCs). These findings indicate that heterogeneous LSC subpopulations with distinct developmental phenotypes can co-reside in the same patient. Moreover, LSC heterogeneity gives rise to bulk tumor populations with differing therapeutic response in PDX models, implying clinical significance.

**Fig. 2.**
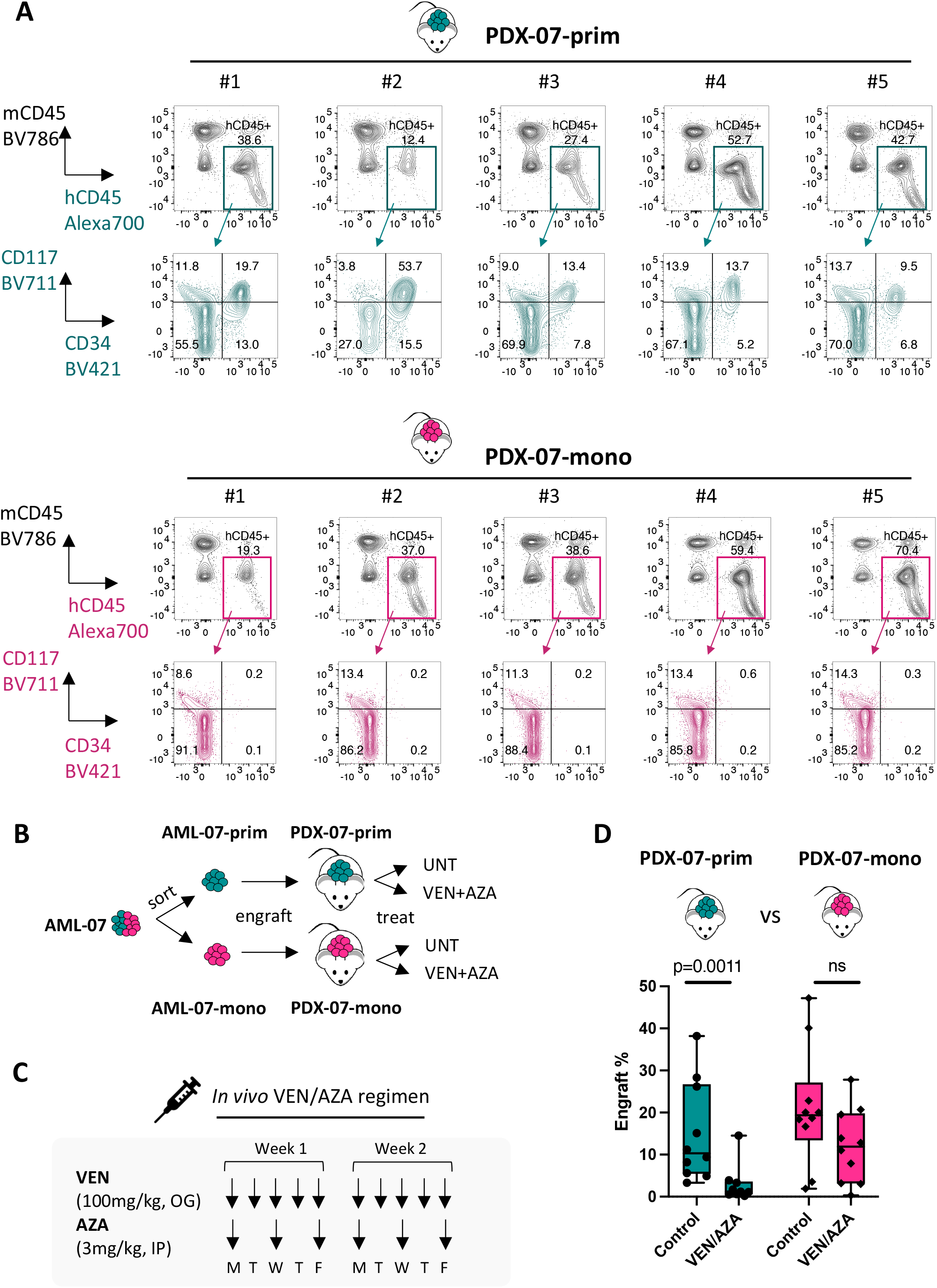
Differing nature of disease arising from prim and mono subpopulations of Multi-MMP AMLs. **A**, Representative flow plots showing immunophenotypic differences between leukemia arising from prim and mono subpopulations of Multi-MMP AML-07 in NSG-S mice. Five representative mice from each group are shown. For each mouse, engrafted human leukemic cells were gated as hCD45+/mCD45- (teal and pink gates) and their expression pattern of CD34 and CD117 were shown to illustrate immunophenotypic differences between the two groups. **B**, A diagram depicting workflow used to isolate primitive and monocytic subpopulations of AML-07 for injecting into PDX mice and subsequent determination of their relative sensitivity to the VEN+AZA regimen in vivo. **C**, Design of the VEN/AZA in vivo regimen (VEN, 100mg/kg, oral gavage (OG), 5 days/week x 2 weeks; AZA, 3mg/kg, Intraperitoneal injection (IP), 3 days/week x 2 weeks). **D**, Impact of *in vivo* VEN+AZA treatments on leukemia engrafted from prim versus mono subpopulations of AML-07. Engraft% was determined by % of hCD45+/mCD45-cells within total viable bone marrow cells. Each dot represents a unique mouse. PDX-07-prim (Control, n=10; VEN/AZA, n=10), PDX-07-mono (Control, n=10; VEN/AZA, n=10). Box plots show median +/- interquartile. Two-tailed Mann-Whitney test is used. ns, not significant.

### Clinical outcomes as a function of m-LSCs

Based on the above findings and our previous studies(12), we hypothesized that de novo AML patients would differ in pathogenesis and clinical outcome as a function of the presence of m-LSCs. As illustrated schematically in **Fig. 3A**, we predict that upon receiving VEN+AZA therapy, Uni-MMP patients that initially present with only p-LSCs should achieve more durable remission due to intrinsic reliance of p-LSCs on venetoclax target BCL-2(12). In contrast, Multi-MMP patients with a distinct m-LSC population are predicted to relapse relatively quickly with monocytic disease. To test this concept, three de novo AML patients who had been treated with the VEN+AZA therapy were evaluated. As illustrated in **Fig. 3B-D**, at diagnosis, patient Pt-20 had no detectable m-LSC activity, whereas patients Pt-12 and Pt-69 had readily detectable m-LSC activities as revealed by xenograft assays. The presence of functionally defined m-LSCs directly correlated with clinical outcomes, where Uni-MMP patient Pt-20 experienced prolonged remission for more than 3.5 years, and Multi-MMP patients Pt-12 and Pt-69 had relatively rapid relapse of disease with a monocytic phenotype within 12 and 3 months, respectively. We subsequently analyzed a cohort of 25 AML patients who relapsed from the VEN+AZA therapy (**Supplementary Table S2)**. We found that nine patients, including five phenotypically monocytic and four who were primitive at diagnosis, relapsed with monocytic features, representing 36% of overall cases **(Fig. 3E**). Importantly, patients who relapsed with monocytic features also had a significantly shorter remission duration than patients who relapsed with a primitive immunophenotype (**Fig. 3F** and **Supplementary Fig. 3A**), suggesting that patients with pre-existing m-LSCs may represent an especially poor prognosis subgroup when receiving VEN+AZA therapy. Overall, these findings indicate that the presence of m-LSCs in newly diagnosed AML patients represents a distinct disease entity and directly predicts pathogenesis of disease in response to VEN+AZA therapy.

**Fig. 3.**
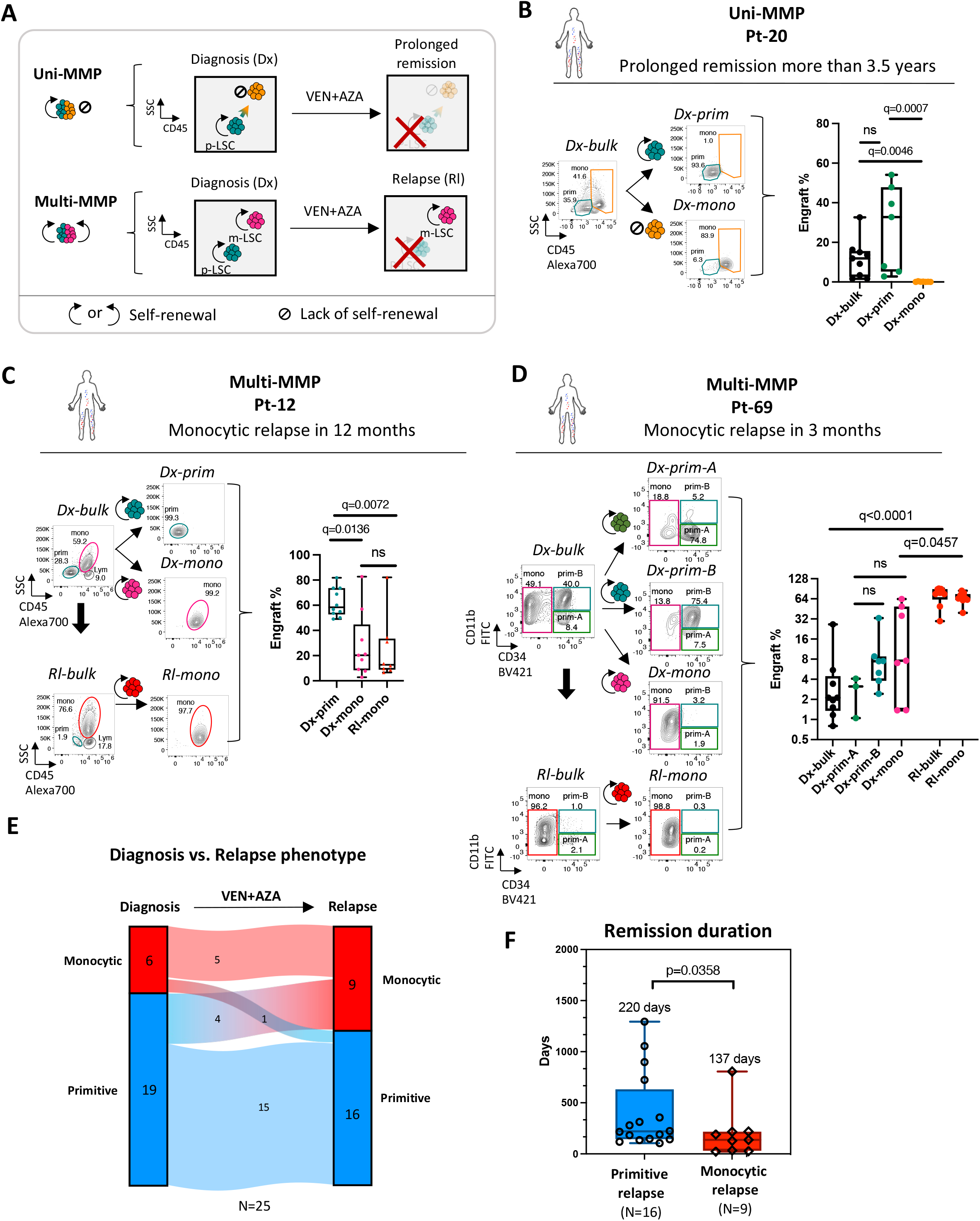
Clinical outcomes as a function of m-LSCs. **A**, A diagram describing the leukemogenesis process of Uni-MMP and Multi-MMP AML patients and their predicted clinical responses to VEN+AZA therapy. **B-D**, Representative cases of Uni-MMP and Multi-MMP AML patients received VEN+AZA therapy. The left panels depict sorting strategies for obtaining diagnosis-prim (Dx-prim) and diagnosis-mono (Dx-mono) subpopulations from diagnosis bulk disease (Dx-bulk), as well as relapse-mono (Rl-mono) subpopulations from relapse bulk disease (Rl-bulk) when applicable. The right panel shows engraft% in NSG-S mice determined by hCD45+/mCD45-% within total viable bone marrow cells. Each dot represents a unique mouse. Median +/- interquartile. Mann-Whitney tests. ns, not significant; *p<0.05, **p<0.01, ***p<0.001. **B**, Patient 20 (Pt-20), a case of Uni-MMP AML presenting prolonged remission after VEN+AZA therapy for more than 3.5 years. Dx-bulk (n=9), Dx-prim (n=7), Dx-mono (n=7). **C**, Patient 12 (Pt-12), a case of Multi-MMP AML presenting predominant monocytic relapse in 12 months after receiving VEN+AZA therapy. Dx-prim (n=10), Dx-mono (n=9), Rl-mono (n=8). **D**, Patient 69 (Pt-69), a case of Multi-MMP AML presenting quick relapse in 3 months post VEN+AZA therapy. In this particular case, prim and mono subpopulations were gated using a different sorting strategy based on primitive antigen CD34 and monocytic antigen CD11b. For the patient’s diagnosis sample, the CD34+/CD11b-, CD34+/CD11b+, and CD34-/CD11b-pp subpopulations were sorted as Dx-prim-A, Dx-prim-B, and Dx-mono subpopulations, respectively. For the patient’s relapse sample, the CD34-/CD11b-pp subpopulation was sorted as the predominant Rl-mono subpopulation. Dx-bulk (n=9), Dx-prim-A (n=3), Dx-prim-B (n=7), Dx-mono (n=7), Rl-bulk (n=9), Rl-mono (n=9). **E**, Phenotypic changes from diagnosis to relapse in a cohort of AML patients received VEN+AZA therapy (N=25, Supplementary Table S2). **F**, Remission duration for AML patients with monocytic relapse (N=9) versus non-monocytic relapse (N=16) (Also see Supplementary Fig 3). Each dot represents a unique patient. Median duration time of both groups are shown in days. In b, c, d, and f, box plots represent median +/- interquartile. In b-d, Kruskal-Wallis test was used. In f, one-tailed Mann-Whitney test is used. ns, not significant.

### Identification of the m-LSC immunophenotype

Given the unique developmental properties and significant therapeutic impact of m-LSCs, we next sought to identify them in primary AML specimens and begin to characterize their biology. To this end, we employed CITE-seq (Cellular Indexing of Transcriptomes and Epitopes by Sequencing) analysis allowing simultaneous measurement of protein-based surface antigens (listed in **Supplementary Table S3**) and RNA-based transcriptome analysis at a single cell level. We performed CITE-seq analysis on a cohort of 27 primary AML specimens containing immunophenotypically defined Prim, MMP, and Mono AMLs (listed in **Supplementary Table S4**). The CITE-seq data was first analyzed using Clustifyr, an application that assigns phenotypes based on comparison to the transcriptome of normal human hematopoiesis(15-17). This analysis classifies myeloid cells into HSPC, MPP, Early Promyelocyte-like, Late promyelocyte-like, Myelocyte-like, classical monocyte-like, and Nonclassical monocyte-like subclusters, and clearly reveals the myeloid developmental spectrum within primary AML specimens (**Fig. 4A**). The lineage assignments are strongly supported by the expression patterns of classic lineage-specific markers such as CD7, CD56, CD19, CD33, CD34, CD11b, and CD123 at both protein and RNA levels (**Supplementary Fig. 4A-B**), as well as lineage-specific transcriptional factors such as RUNX1, SPI1, TCF7, PAX5 etc. (**Supplementary Fig. 4C**). Next, the overall CITE-seq data were segregated into Prim, MMP, and Mono groups according to their immunophenotype. We also further divided the MMP group into Uni-MMP and Multi-MMP subgroups based on their functional LSC activities measured by xenograft assay (**Fig. 4C**). The relative proportion of each myeloid subcluster within Prim, Uni-MMP, Multi-MMP, and Mono groups was highly concordant with their expected identity. The Prim group contains the highest proportion of HSPC, MPP, Early Promyelocyte-like cells, while the Mono group contains the greatest frequency of Myelocyte/Classical/Nonclassical monocyte-like cells, and the MMPs contain intermediate level of both (**Fig. 4B**).

**Fig. 4.**
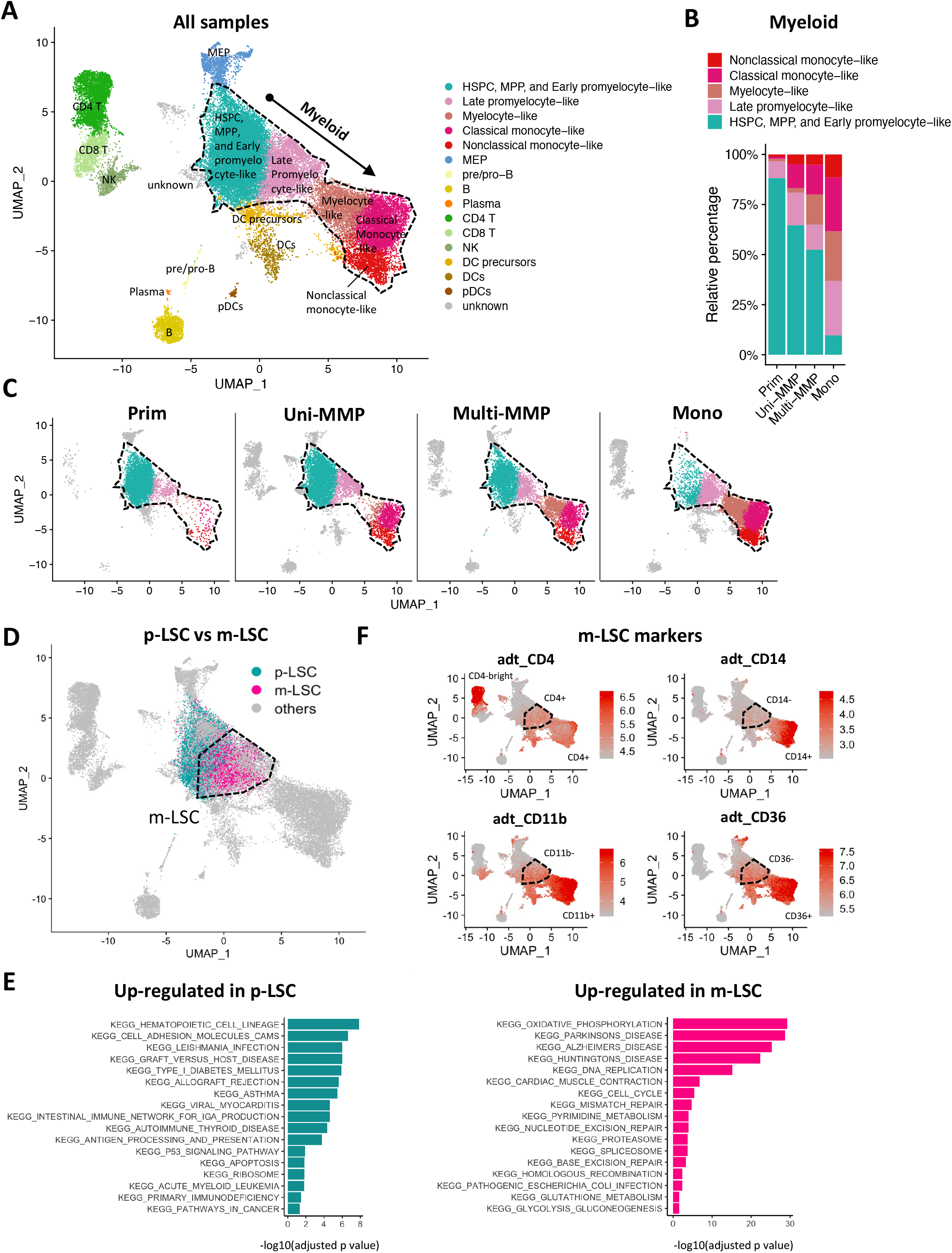
Identification of the m-LSC immunophenotype. **A**, A UMAP of primary AML specimens (N=27) containing cells of myeloid, lymphoid, and erythroid lineages. Information of specimens are detailed in Supplementary Table S4. Dotted lines highlight myeloid disease relevant subclusters. **B**, Stacking bar graphs showing relative proportion of each subcluster within the highlighted “myeloid” region of the UMAP. **C**, UMAPs of immunophenotypically determined Prim, Mono, and MMP AMLs. The MMP AMLs were further split into Uni-MMP and Multi-MMP subgroups based functional xenograft assays as described in Fig. 1. Grouping information are detailed in Supplementary Table S4. **D**, Assignment of p-LSC and m-LSCs based scoring of p-LSC and m-LSC specific transcriptional signatures. **E**, Top up-regulated pathways in p-LSCs and m-LSCs relative to each other as determined by GSEA analysis. **F**, Protein expression of surface antigens CD45, CD34, CD4, CD14, CD11b, and CD36. The m-LSC enriched region is highlighted by dotted lines.

To begin to identify the m-LSC immunophenotype based on CITE-seq data, we first sought to generate a m-LSC specific gene expression signature. To this end, we reasoned that the gene expression pattern of KMT2A-rearranged leukemia is a good approximation of m-LSC biology since studies have shown that the KMT2A-rearranged leukemia is distinct from other AMLs(18), frequently associated with a monocytic immunophenotype(19), and has LSC characteristics that are distinct from conventional CD34+ primitive LSCs(20). Therefore, we generated a candidate m-LSC-specific gene expression signature through combining several KMT2A-rearranged LSC signatures by Hess et al and Somervaille et al (21-23). In parallel, we also generated a p-LSC-specific signature based on studies of conventional CD34+ primitive LSCs by Eppert et al and Ng et al (24, 25). Through scoring the custom m-LSC and p-LSC specific signatures (**Supplementary Table S5**) on each myeloid cell, we found that the p-LSCs and m-LSCs are largely located within the HSPC, MPP, Early Promyelocyte-like and the Late promyelocyte-like subclusters, respectively (**Fig. 4D** and **Supplementary Fig. 4D**). We subsequently identified significantly up-regulated genes in both types of LSCs and performed GSEA analysis (**Supplementary Table S6 and S7**). These studies showed that m-LSCs are uniquely enriched for pathways regulating oxidative phosphorylation, DNA replication, cell cycle, and glutathione metabolism, suggesting more active metabolism, cell cycle status, and redox homeostasis (**Fig. 4E, Supplementary Fig. 4E**, and **Supplementary Table S8**). More importantly, expression analysis of the surface antigens revealed that transcriptionally defined m-LSCs are largely CD4+, CD14-, CD11b-, and CD36-, relative to other myeloid leukemia cells, providing a candidate immunophenotype for m-LSCs (**Fig. 4F**).

### Functional validation of m-LSCs

Based on the CITE-seq analysis in **Fig 4**, we next performed a series of flow sorting and transplant studies to functionally validate the m-LSC immunophenotype. We initially employed two mono specimens AML-16 and AML-20, and isolated various subpopulations of cells using expression of CD45, CD34, CD4, CD14, CD11b, and CD36 (**Fig. 5A-B** and **Supplementary Fig. 5A-B**). To avoid contamination from conventional CD34+ p-LSCs, we gated on the CD34-fraction when validating the m-LSCs. As shown in **Fig. 5D-E** and **Supplementary Fig. 5D-E**, functional analysis of each subpopulation demonstrated that the majority of m-LSCs are enriched in the immunophenotype of CD34-, CD4+, CD14-, CD11b-, and CD36-. Importantly, secondary transplantation of animals originally engrafted with m-LSCs (designated as population D) also demonstrated robust engraftment, an indication of strong self-renewal potential. Notably, in AML-20, some engraftment potential was detected in the CD14-, CD11b+, and CD36+ counterparts (**Fig. 5E**), suggesting that in certain patients the developmental hierarchy of m-LSC-driven AMLs can be shallower than others (e.g., AML-16). We also applied a similar approach to validate m-LSCs in Multi-MMP patients. To this end, we employed AML-07 that was functionally defined to be a Multi-MMP specimen (**Figs. 1-2**). As shown in **Fig. 5C, 5F** and **Supplementary Fig. 5C, 5F**, serial transplant experiments demonstrated that m-LSCs in this Multi-MMP AML were also exclusively enriched by the CD34-, CD14-, CD11b-, and CD36-immunophenotype. Of note, CD4 was not included in this sort due to limited cell numbers. Together, these data reveal a m-LSC immunophenotype that is applicable to both Mono and Multi-MMP AMLs, entirely distinct from profiles previously described for the more conventional primitive LSCs (i.e., CD34+, CD38-, etc.) (4, 24, 25).

**Fig. 5.**
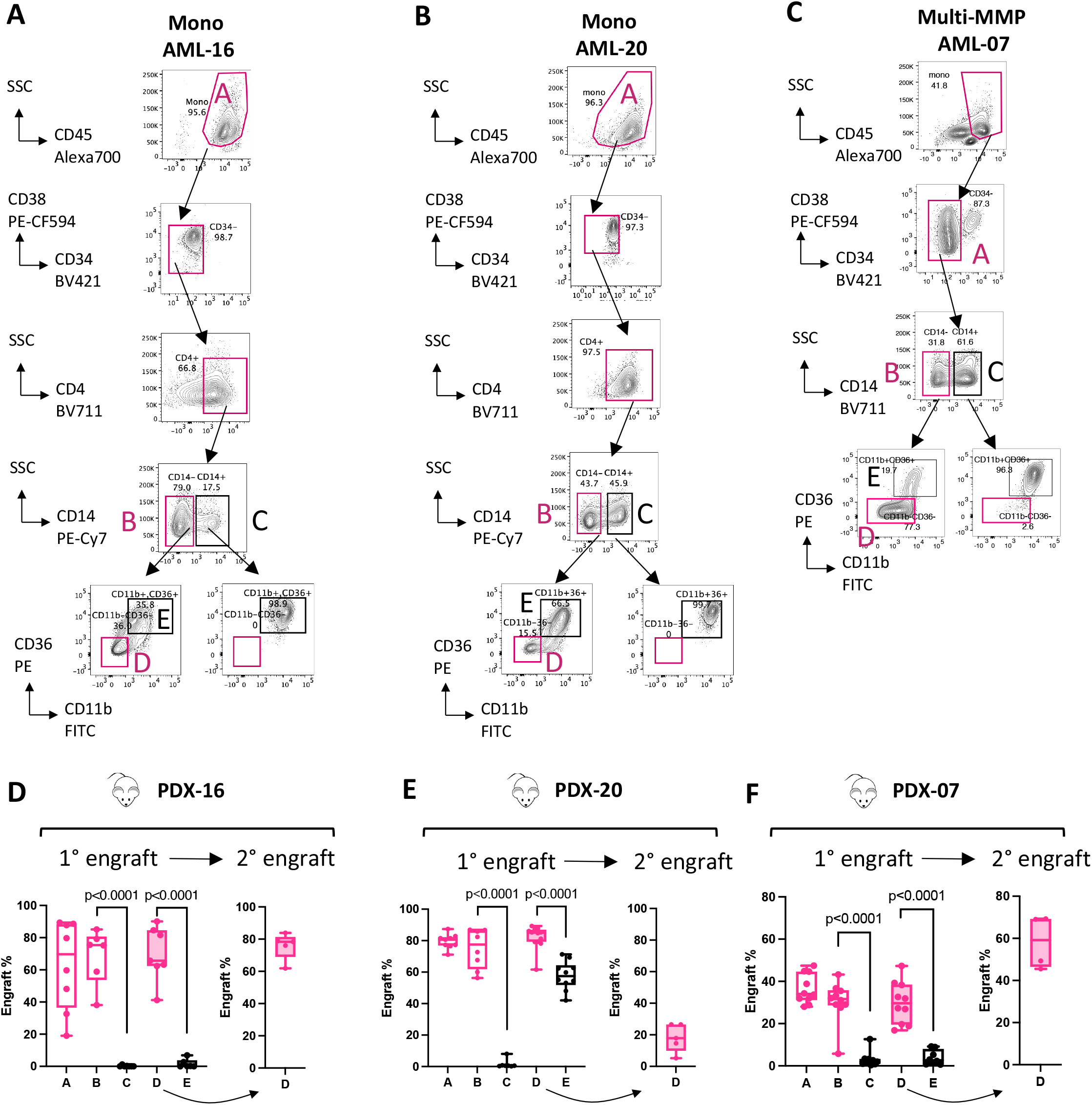
Functional validation of m-LSC immunophenotypes. **A-C**, Gating strategies for sorting various subpopulations of Mono AML-16, Mono AML-20, and Multi-MMP AML-07 for determining their m-LSC activities using xenograft studies. The sorting is detailed in Supplementary Fig. 5a-c. Briefly, for AML-16 and AML-20, A (Live/mono), B (Live/mono/CD34-/CD4+/CD14-), C (Live/mono/CD34-/CD4+/CD14+), D (Live/mono/CD34-/CD4+/CD14-/CD11b-CD36-), and E (Live/mono/CD34-/CD4+/CD14-/CD11b+CD36+) were sorted and engrafted. For AML-07, CD4 was not included in the sort due to limitation of cells. **E-F**, Results from transplanting subpopulations of AML-16 (A(n=8), B(n=6), C(n=7), D(n=7), E(n=6)), AML-20 (A(n=9), B(n=8), C(n=7), D(n=12), E(n=10)), AML-07 (A(n=10), B(n=10), C(n=9), D(n=10), E(n=8)) are shown as PDX-16, PDX-20, and PDX-07, respectively. Engraft% was determined by % of hCD45+/mCD45-cells within total viable bone marrow cells. Each dot represents a unique mouse. Box plots represent median +/- interquartile. Two-tailed Mann-Whitney tests.

To further explore the immunophenotype of m-LSCs, we also used analytical flow cytometry to evaluate the expression of several additional cell surface antigens associated with stem cell activity. As previously reported by Quek et al, both CD117 and CD244, were found to be frequently expressed specifically in CD34-AML specimens (8). As shown in **Supplementary Fig 5G**, there was no detectable expression of CD117 in any of the seven specimens that underwent functional evaluation for the presence of m-LSCs. Similarly, CD244 was only detected at low levels in two of seven specimens. We also investigated expression of the monocytic marker, CD64, as this antigen was prevalent in our original characterization of monocytic relapse (12). We found that CD64 was strongly expressed in six of seven specimens. Thus, while we do not yet have functional validation of these antigens, we propose that the immunophenotype of m-LSCs will be CD117-, CD244-, and CD64+.

### Targeting purine metabolism in m-LSCs

Finally, given the significance of m-LSCs in driving refractory/relapse response to venetoclax-based regimens, we sought to identify potential m-LSC selective therapies. Using the CITE-seq data described in **Fig. 4**, we determined that several gene expression signatures were evident in the m-LSC compartment including pyrimidine metabolism and purine metabolism (**Fig. 6A-B**). These results were corroborated by expression patterns of key enzymes known to modulate one-carbon metabolism (e.g., TYMS and DHFR) that feed into purine (e.g., HPRT1) and pyrimidine synthesis (e.g., DHODH) pathways (**Fig. 6C**). Hence, we performed functional studies to evaluate agents known to impact these pathways. Specifically, we chose methotrexate (MTX), brequinar (BRQ), and cladribine (CdA), based on previous reports of their respective activities in inhibiting one-carbon metabolism enzymes DHFR and TYMS (26), pyrimidine synthesis enzyme DHODH (27), and purine-based DNA/RNA synthesis (28). Colony-forming Unit (CFU) assays were performed on m-LSCs isolated from AML-16 and AML-20, along with normal CD34+ hematopoietic stem and progenitor cells (HSPC) from two healthy donors. As shown in **Fig. 6D**, all three agents showed AML-specific toxicity at doses that were negligibly toxic to normal HSPC controls. The purine analogue CdA was particularly effective in eradicating the stem and progenitor potential of m-LSCs. Thus, *in vivo* proof-of-concept studies were performed using CdA in combination with VEN+AZA. As shown in **Figs. 6E** and **Supplementary Fig. 6A**, MMP AML-07 was transplanted into a cohort of NSG-S mice and treated with CdA alone, VEN+AZA alone, or the combination. Analysis of AML cells in both bone marrow and spleen clearly demonstrate enhanced *in vivo* targeting with the combined regimen (**Fig. 6F and Supplementary Figs. 6B-D**).

**Fig. 6.**
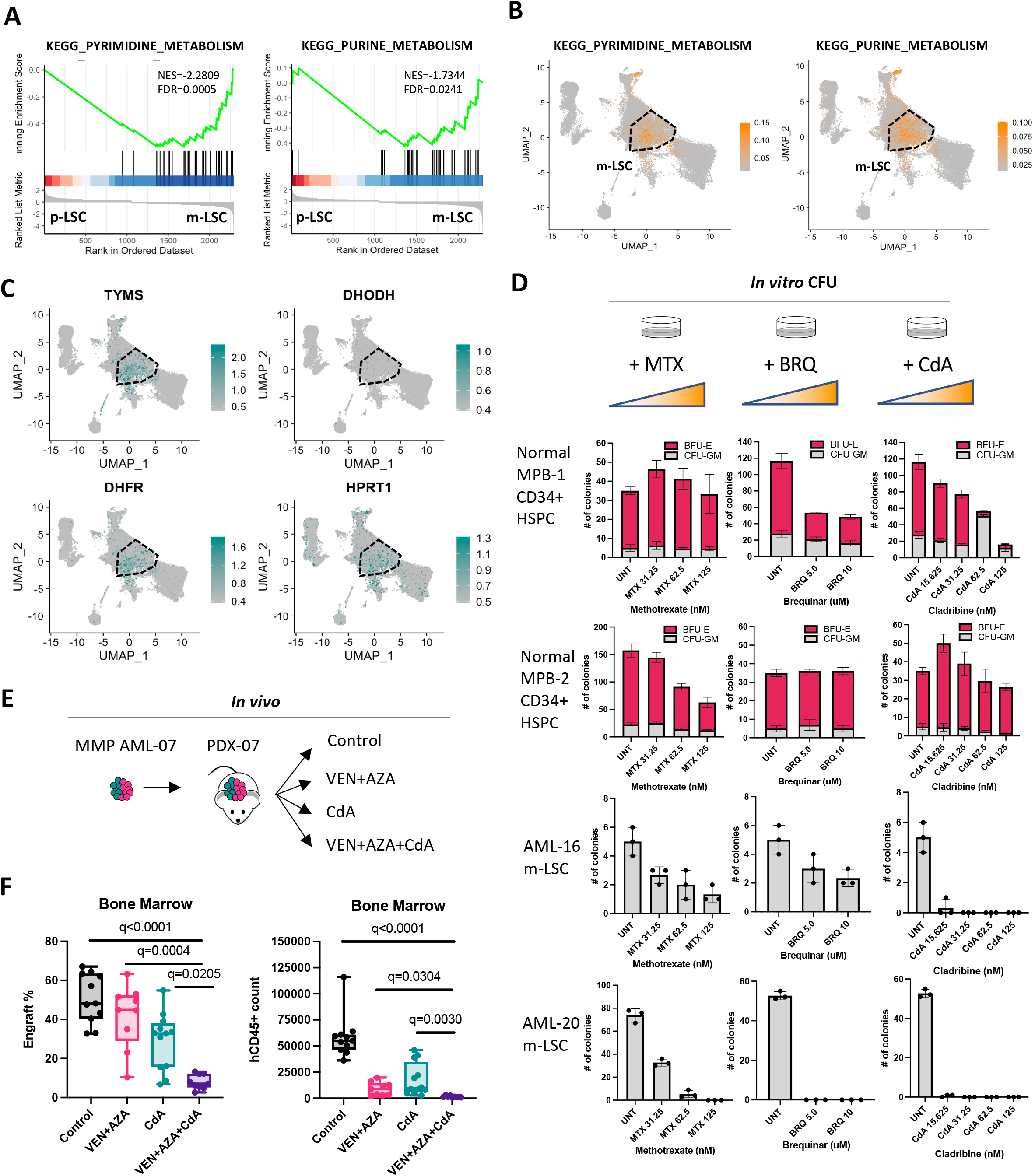
Molecular properties and targeting of m-LSCs. **A**, GSEA enrichment plots showing enrichment of pyrimidine and purine metabolism pathways in m-LSCs comparing to p-LSCs. **B-C**, Scoring of pyrimidine and purine metabolism pathways and expression of TYMS, DHFR, HPRT1, and DHODH on UMAP. The dotted lines highlight the m-LSC enriched region. **D**, Impact of TYMS inhibitor methotrexate (MTX), DHODH inhibitor brequinar (BRQ), and purine analogue cladribine (CdA) on colony-forming unit (CFU) potential of CD34+ HSPCs isolated from two normal mobilized peripheral blood samples (MPB-1, 2) versus mono-LSCs isolated from mono AML-20 and mono AML-16. **E**, A diagram depicting workflow used to measure in vivo efficacy of cladribine (CdA) in combination with VEN+AZA in treating a PDX model of Multi-MMP AML. VEN, 100mg/kg, OG, 5 days/week x 2 weeks; AZA, 3mg/kg, IP, 3 days/week x 2 weeks, CdA, 10mg/kg, IP, 3 days/week x 2 weeks. **F**, Impact of VEN+AZA, CdA or combo treatments on the bone marrow tumor burden of PDX. Engraft% was determined by % of hCD45+/mCD45- cells within total viable bone marrow mononuclear cells. hCD45+ count was determined by direct quantification of hCD45+/mCD45-cells within a set volume of marrow harvest using flowcytometry. Each dot represents a unique mouse. Control (n=11), VEN+AZA (n=9), CdA (n=12), VEN+AZA+CdA (n=9). Box plots represent median +/- interquartile. Kruskal-Wallis test was used. ns, not significant.

## Discussion

The findings presented in this report describe the identification and characterization of a previously unrecognized type of human AML stem cell that we term m-LSC. This particular subclass of LSC is distinguished from more primitive subtypes by virtue of a unique immunophenotype (CD34-, CD4+, CD14-, CD11b-, and CD36-), a relatively narrow developmental profile that is limited to the creation of monocytic progeny, and a gene expression profile that is roughly analogous to normal human promyelocytes. Notably, the m-LSC is distinct from the CD34-LSC populations described previously by Quek et al, where expression of both CD177 and CD244 were prevalent (8). The molecular biology of m-LSCs differs from more primitive LSCs in that BCL-2 dependency seems to be largely dispensable, making this type of LSC resistant to treatment with venetoclax and azacitidine (29). Further, m-LSCs demonstrate selective reliance on one-carbon metabolism and purine/pyrimidine metabolism, adding to the importance of cellular metabolism in the context of AML pathogenesis and therapeutic resistance (30-34). Importantly, reliance on purine metabolism appears to mediate increased sensitivity to agents such as cladribine, a well-known purine analogue. We demonstrate that addition of cladribine to the VEN+AZA regimen increases eradication of primary AML containing m-LSC activity in both *in vitro* and *in vivo* preclinical models. We believe these findings provide important mechanistic insights that may explain results from several recent clinical trials reporting superior CR/CRi and MRD negativity rates when venetoclax is combined with chemotherapy regimens especially ones that contain cladribine or its close analogue fludarabine, namely FLAG-Ida, CLIA or Clad-LDAC/AZA (35-38). Notably, the addition of cladribine in these three trials was mainly based on empirical clinical experience and the long history of using cladribine-containing chemotherapy regimens for relapse/refractory stages of AML. Our discovery of m-LSCs and their unique resistance to venetoclax and sensitivity to cladribine suggests that the efficacy of cladribine in combination with venetoclax-based regimens may be at least in part due to more comprehensive eradication of the overall LSC population.

The collective findings from our work suggest stratification of primary AML patients based on the nature of underlying LSC subpopulations. As shown in **Fig. 7**, we use the tree as an analogy to describe developmental hierarchy of AML where the underground roots represent LSCs and the branches above the ground symbolize more differentiated blasts. In **Fig. 7A**, A single class of more primitive LSCs (p-LSCs) may be present in newly diagnosed AML patients, with varying degrees of monocytic differentiation potential. In this scenario, no m-LSCs are present, and venetoclax-based therapy would be predicted to confer relatively high CR rates and longer remissions. In contrast, as shown in **Fig. 7B**, some newly diagnosed Multi-MMP AML patients present with at least two distinct subtypes of LSC with primitive vs. monocytic characteristics (p-LSC and m-LSC). Depending on the size of the m-LSC population, these patients would clinically be expected to respond and then relapse, or to be refractory to venetoclax-based therapies. Finally, in more extreme cases, only m-LSCs are present (**Fig. 7C**), which would likely result in disease that is refractory to venetoclax. Notably, this is exactly what we previously reported: as shown in **Supplementary Fig. 7A-B**, patients that present with the most differentiated monocytic phenotypes (i.e. FAB-M5) but not those with myelomonocytic phenotypes (i.e. FAB-M4) show the highest frequency of refractory disease (12, 13). Moreover, as shown in **Fig. 3F**, patients relapsed with a monocytic disease have significantly shorter duration of remission comparing to ones that had primitive relapse.

**Fig. 7.**
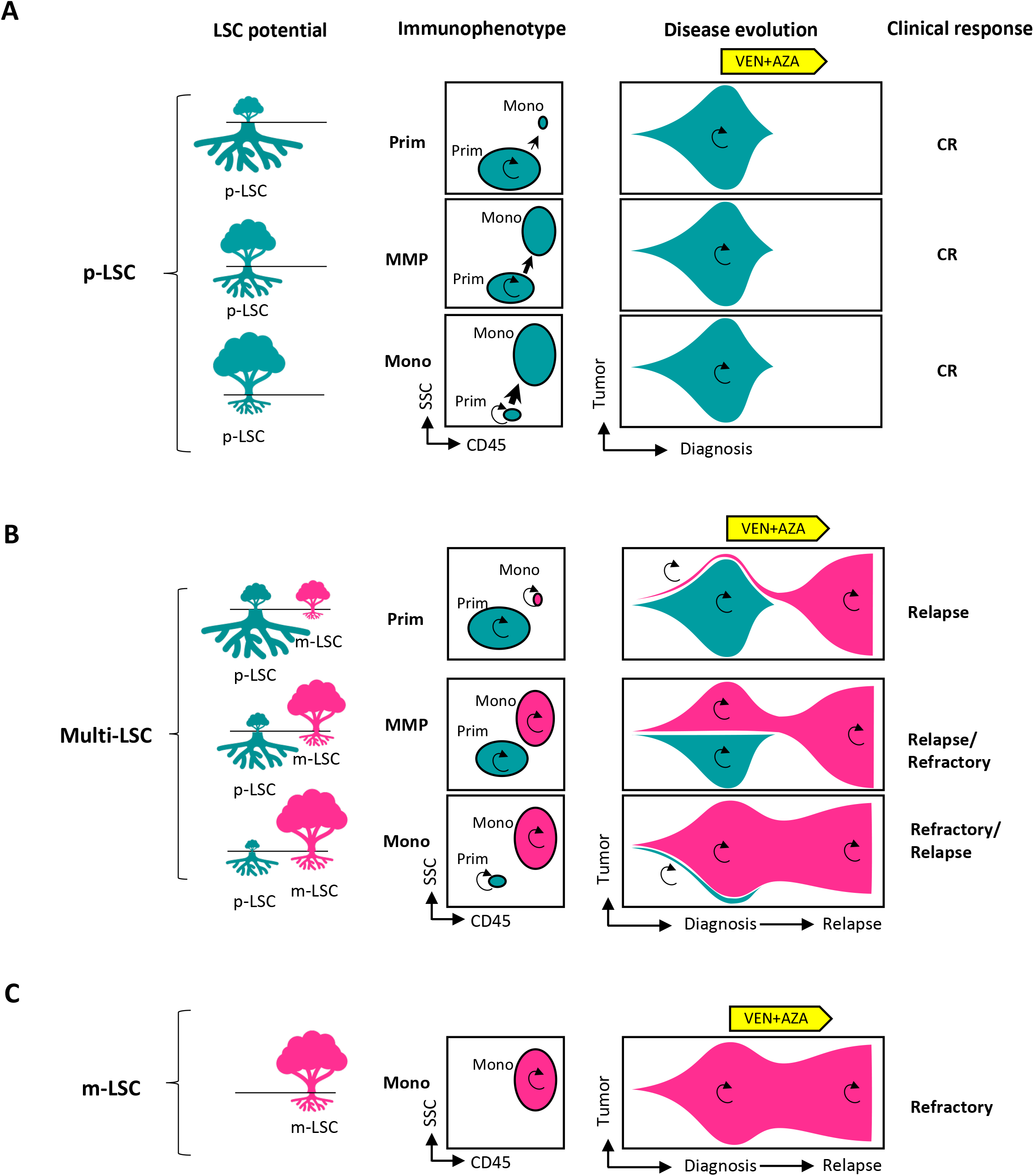
A model depicting interplay between LSC potential, immunophenotype, disease evolution, and clinical response to the venetoclax plus azacitidine therapy in AML patients. **A**, A first group of AML patients with disease solely driven by p-LSCs captured at various maturation stages with predominant Prim, MMP, and predominant Mono immunophenotypes. **B**, A second group of AML patients with multi-LSC activities (contains both p-LSCs and m-LSCs) presenting Prim, MMP, or Mono immunophenotypes depending on the relative degrees of maturation and ratio of the two diseases rooted from the two distinct LSC subtypes. **C**, A last group of AML patients with disease solely driven by m-LSCs usually presenting a predominant mono immunophenotype due to the inherent nature of m-LSCs that are already resting at a relatively more mature promyelocyte-like developmental stage. In all graphs, teal colored symbols represent p-LSCs and their progenies, pink colored symbols represent m-LSCs and their progenies, circular arrows represent self-renewal capacity.

The model described in **Fig. 7** has specific ramifications for the design of future therapies. Detection of any m-LSC population at diagnosis in a patient who is being considered for a venetoclax-based therapy may warrant consideration of additional therapies designed to selectively target monocytic population, in the hopes that relapse, which carries a very poor prognosis in this setting (39, 40), can be avoided. Aside from cladribine (and methotrexate, **Fig. 6**), such potential agents include immunotherapies directed towards monocytic antigens such as CD64 and LILRB4, or small molecules that selectively impair unique biology of phenotypically monocytic AML. For example, as we and others have previously reported, MCL1 inhibitor drugs appear to be more active in the context of monocytic AML (12, 41). Our data suggest this is due to dependence on MCL1 for energy metabolism in m-LSCs (12). Alternatively, for patients with pure p-LSC disease at diagnosis, no additional therapy beyond venetoclax and azacitidine may be appropriate to allow for durable remissions, and for those with pure m-LSCs, one could consider replacement of venetoclax for another therapeutic strategy more likely to be effective. Future clinical trials designed in this way may serve to optimize outcomes and prevent over-treatment.

Importantly, m-LSC driven disease propagation and/or relapse is only observed in approximately 30% of AML patients. However, the emergence of drug resistant disease eventually occurs in the majority of newly diagnosed patients treated with the VEN+AZA regimen. This finding strongly implies that other subclasses of LSCs are present (or can evolve) in patients treated with venetoclax-based regimens. Thus, defining additional phenotypes associated with venetoclax-resistant LSCs remains a high priority.

Finally, our study serves to further elaborate the connection between disease-initiating mutations, LSC heterogeneity, developmental state of AML, and varying therapeutic response in the clinic. Previous studies have established several key elements of this broader picture. First, a recent single cell multi-omic study by Miles et al. has drawn strong ties between mutational profiles and immunophenotypes of AML (10). The study found venetoclax-sensitive IDH mutants tend to express the primitive marker CD34 and conversely venetoclax-resistant RAS mutants usually express monocytic antigen CD11b (10, 42). Second, deconvolution analysis of bulk RNAseq data from approximately 1,000 primary AML patients by Zeng et al has demonstrated a similar strong association between genotype, phenotype, and drug response at bulk level (14). Of note, a link between RAS mutations and mature monocytic differentiation was particularly strong in this study (14). Third, a recent human study by Zeisig et al. showed that LSC activities exclusively exist in the CD34-but not the conventional CD34+ subfraction of KMT2A-rearranged primary AML displaying either CD34-/CD34low or CD34+ immunophenotypes (43). Lastly our previous study suggests RAS-driven monocytic subclones can be a major driver of monocytic relapse in AML patients treated with VEN+AZA. Our current study shows that a novel type of venetoclax-resistant m-LSC can be found in both RAS-mutant mono (AML-20), KMT2A-rearranged mono (AML-16), and Multi-MMP (AML-07) contexts (**Fig. 5**). Thus, taken together, these lines of investigation suggest that complex oncogenic mutations likely converge on limited LSC subtypes that can shape the developmental hierarchy of bulk disease and ultimately affect therapeutic response in the clinic. This model implies that understanding LSC heterogeneity is at the core of improving therapeutic interventions for AML.

Finally, besides AML, a recent study found that stem cell architecture can also impact disease progression and predict venetoclax response in myelodysplastic syndrome (44), suggesting malignant stem cell heterogeneity has broad ramifications in myeloid pathogenesis. Thus, we postulate that successful development of the next generation of precision medicine towards AML requires careful dissection of human LSC heterogeneity and discovery of targeted therapies tailored to each LSC subtype.

## Methods

### Primary AML and normal mobilized peripheral blood samples

Primary human AML specimens were obtained from leukapheresis product, peripheral blood, or bone marrow of AML patients who gave informed consent for sample procurement on the University of Colorado tissue procurement protocols (Colorado Multiple Institutional Review Board Protocol #12-0173 & #06-0720). Normal mobilized peripheral blood (MPB) specimens were obtained from volunteer donors at the University of Colorado. Age, sex, cytogenetics, and mutation information of primary AML specimens used in the current study are detailed in **Supplementary Table S1**.

### Patients, Treatment and Responses

Twenty-five newly diagnosed AML patients who received venetoclax + azacitidine therapy and had experienced relapse response were included in this study. The patients were diagnosed, treated, and monitored for relapse response over a period from January 2015 to February 2020. The University of Colorado Institutional Review Board approved a request to retrospectively analyze these patients (#19-0115). Diagnosis and relapse phenotypes were determined through morphological review by hematologists and examination of clinical flow notes when available. Remission duration was calculated as the numbers of days in between the date of best response and the date of relapse. Cytogenetic, mutational information of diagnosis and relapse stages as well as remission duration time are detailed in **Supplementary Table S2**.

### Processing and culturing of primary AML and normal CD34+ HSPCs

Primary human AML specimens and normal MPB samples were resuspended at about 100-200 e6 cells per ml in freezing media composed of 50% FBS (GE Healthcare), 10% DMSO (Sigma) and 40% IMDM media (GIBCO) and then cryo-preserved in liquid nitrogen. Cells were thawed in 37°C water bath, washed twice in thawing media composed of IMDM (GIBCO), 2.5% FBS (GE Healthcare) and 10 ug/ml DNase (Sigma). Normal CD34+ HSPCs were enriched from thawed MPB samples using the CD34 MicroBead kit (Miltenyi Biotec). Cells were cultured in complete serum-free media (SFM) in 37°C, 5% CO2 incubator. SFM is composed of IMDM (GIBCO), 20% BIT 9500 (STEMCELL Technologies), 10ug/ml LDL (Low Density Lipoprotein, Millipore), 55uM 2-Mercaptoethanol (GIBCO) and 1% Pen/Strep (GIBCO). Complete SFM were made by supplementing the SFM with FLT-3, IL-3 and SCF cytokines (PeproTech), each at 10 ng/ml.

### Colony-forming assays

Freshly sorted m-LSCs from primary AMLs or prepared CD34+ HSPCs isolated from normal mobilized peripheral samples were plated in human methylcellulose (R&D systems) at about 100K/ml and 2K/ml, respectively. Small molecule inhibitors were directly added into the methylcellulose at the desired final concentration at the plating. Colonies were counted 2-3 weeks after the initial plating.

### Immunophenotyping of primary AML

About 0.5-1e6 freshly thawed primary AML cells were stained with immunophenotyping panel containing antibodies against human CD45, CD34, CD117, CD11b, CD64, CD14, CD244, LILRRB4, CD36 or CD68 at 4°C for 15 mins, washed with ice cold FACS buffer, and resuspended in FACS buffer and analyzed on BD FACS Celesta flowcytometry (BD). FCS files were analyzed on Flowjo 10.5.3 (Flowjo). A complete list of flow antibodies is listed in **Supplementary Table S3**.

### Flow sorting of prim and mono subpopulations for xenograft studies

Freshly thawed primary AML cells were stained with viability dyes and antibodies against human CD45 or CD34 and CD11b. For all specimens except for Pt-69, the prim subpopulation was sorted as CD45-medium, SSC-medium, while the mono subpopulation was sorted as CD45-bright, SSC-medium/high as described previously(12). For Pt-69, prim and mono subpopulations were not readily separable using the CD45/SSC gating strategy. An alternative CD34/CD11b gating strategy was used. From the CD34/CD11b gate, two prim subpopulations were revealed from the diagnosis (Dx) specimen, one as Dx-prim-A displaying a CD34+/CD11b-phenotype, one as Dx-prim-B bearing a CD34+/CD11b+ phenotype. One mono subpopulation was also revealed from the diagnosis (Dx) specimen, named as Dx-mono exhibiting a CD34-/CD11b-partial positive (PP) phenotype. In contrast, at relapse (Rl), a single mono subpopulation showing a CD34-/CD11b-PP phenotype was revealed as Rl-mono. All subpopulations were sorted and used for xenograft studies.

### Xenograft studies

NSG-S (NOD.Cg-Prkdcscid Il2rgtm1Wjl Tg(CMV-IL3,CSF2,KITLG)1Eav/MloySzJ) mice (The Jackson Laboratory) were used for xenograft studies in this study as previously described(45). Male or female mice ranging in age from 6 to 8 weeks were started on experiment. Littermates of the same sex were randomly assigned to experimental groups. NSG-S mice were pre-conditioned 24 hours prior to transplant with 30mg/kg busulfan (Alfa Aesar) via intraperitoneal (IP) injection. The busulfan stock was made fresh at 25 mg/ml in 100% DMSO, then the stock was diluted 1:10 in pre-warmed saline (0.9% NaCl) down to 2.5 mg/ml right before use. The diluted busulfan solution was kept in 37°C water bath before IP injection to prevent precipitation of busulfan due to low solubility. For comparing the engraftment potential of different subpopulations of a primary AML, each subpopulation was sorted according to their percentage of total viable cells (detailed in **Supplementary Table S9**). When the cell dose was less than 0.5e6/mouse, mononuclear cells isolated from bone marrow and spleens of naïve NSG-S mice were used as carrier cells. The sorted cells with or without addition of carrier cells were then washed, pelleted, and resuspended in saline buffer to allow tail vein injection into NSG-S mice at 0.1ml per mouse. Fifteen minutes prior to injection, in vivo anti-human CD3 antibody (BioCell) was added at a final concentration of 1ug/e6 cells to prevent potential graft versus host disease. During all experiments, the weight of mice was approximately 20-30 grams with no animals losing greater than 10% body weight. The mice were kept in ventilated cages and given in vivo treatments when needed in the vivarium at University of Colorado. Majority of the experiments lasted for 6 to 12 weeks. At the end of the experiments, mice were euthanized using carbon dioxide. Bone marrow and spleen were harvested, subjected to red blood cell lysis, and the mononuclear cells were stained with human CD45, mouse CD45 antibodies, and DAPI to determine percentage of engraftment within viable cells. All animal work were performed in accordance with Institutional Animal Care and Use Committee protocol number 00308.

### In vivo treatments

About 2-4 weeks post initial transplant, tumor burden in the bone marrow was determined to be above 5% in sentinel mice. Mice were then treated with various in vivo regimens as follows. Venetoclax was given at 100mg/kg via oral gavage, five days/week for two weeks; Azacitidine was given at 3mg/kg via intraperitoneal injection, three days/week for two weeks; Cladribine was given at 10mg/kg via intraperitoneal injection, three days/week for two weeks. All treatments were given in the same two-week time window when stated together.

### CITE-seq sample preparation and library construction

Mononuclear cell suspensions were prepared from freshly thawed primary AML specimens cryopreserved in liquid nitrogen. For each specimen, about 1-2e6 mononuclear cell suspension was stained with a panel of Total-seq B antibodies (1ug/each, Biolegend) and fluorochrome conjugated flow antibodies for CD45 (BD Biosciences), CD235a (BD Biosciences), and DAPI. Cells were stained in staining buffer (PBS + 2% FBS) for 20 minutes at 4°C. Viable and red blood cell excluded cells were obtained through sorting DAPI-/CD235a-cells on BD Biosciences ARIA II cell sorter. After sorting, cells were washed twice using staining buffer and resuspended to 1000 cells/μl in PBS with 2% FBS and used immediately for capture. Following cell collection and counting, cells were processed with the 10x Genomics 3’ dual index v3.1 library kit with feature barcoding technology for cell surface proteins. Briefly 10,000 cells are targeted from stock suspension of 1000 cells/μl. Cells were processed following the protocol to generate 3’ gene expression libraries as well as Feature Barcode cell surface libraries. Both libraries were dual indexed, and samples were quantified by Qubit (Life Technologies) and assessed for size and quality by Tapestation (Agilent). Libraries were normalized and pooled for sequencing on a Novaseq 6000 (Illumina) for paired-end 2×150 bp sequencing. Targeted read depth for gene expression libraries was 100,000 reads/cell or ~ 500 million paired-end reads/library. Targeted read depth for cell surface libraries was 40,000 reads/cell or 200 million paired end reads/library. The panel of CITE-seq antibodies are detailed in **Supplementary Table S3**.

### CITE-seq data pre-processing

Raw sequencing data for gene expression, antibody derived tag (ADT; surface protein), and hashing libraries were processed using STARsolo 2.7.8a (46) (https://github.com/alexdobin/STAR/blob/master/docs/STARsolo.md) with the 10X Genomics GRCh38/GENCODE v32 genome and transcriptome reference (version GRCh38_2020A; https://support.10xgenomics.com/single-cell-gene-expression/software/release-notes/build#GRCh38_2020A) or a TotalSeq barcode reference, as appropriate. Hashed samples were demultiplexed using GMM-Demux (47) (https://github.com/CHPGenetics/GMM-Demux). Next, cell-containing droplets were identified using dropkick 1.2.6 (48) (https://github.com/KenLauLab/dropkick) using manual thresholds when automatic thresholding failed, ambient RNA was removed using DecontX 1.12 (49) (https://github.com/campbio/celda) and cells estimated to contain >50% ambient RNA were removed, and doublets were identified using DoubletFinder 2.0.3 (50) (https://github.com/chris-mcginnis-ucsf/DoubletFinder) and removed. Remaining cells were then filtered to retain only those with > 200 genes, 500-80,000 UMIs, < 10-20% of UMIs from genes encoded by the mitochondrial genome (sample dependent based on UMI distributions), < 5% of UMIs derived from HBB, < 20,000 UMIs from antibody-derived tags (ADTs), and >100-2,750 UMIs from antibody derived tags (sample dependent based on UMI distribution). Filtered cells were modeled in latent space using TotalVI 0.18.0 (51) (https://github.com/scverse/scvi-tools) to create a joint embedding derived from both RNA and ADT expression data, corrected for batch effects, mitochondrial proportion, and cell cycle. Scanpy 1.8.2 (52) (https://github.com/scverse/scanpy) was used to cluster the data in latent space using the leiden algorithm(53) and marker genes were identified in latent space using TotalVI.

### CITE-seq data analysis

Clusters were annotated using clustifyr 1.9.1 (15) (https://github.com/rnabioco/clustifyr) and the leukemic/normal bone marrow reference dataset generated by Triana et al (17). Scanpy and Seurat 4.1.1 (54) (https://github.com/satijalab/seurat) were then used to generate UMAP projections from the TotalVI embeddings and perform exploratory analysis, data visualization, etc. p-LSCs and m-LSCs were identified through scoring each individual cell using the AddModuleScore() function of the Seurat software and custom generated candidate p-LSC and m-LSC gene expression signatures as stated in the main text (**Supplementary Table S5**). P-LSC and m-LSC markers were identified using FindMarkers(), GSEA analysis was performed using the GSEA() function of the clusterProfiler software.

### Statistical analysis

Statistical analyses were performed in Prism 9.3.1. Median+/- interquartile range was used to describe summary statistics. One-tailed or two-tailed Mann-Whitney tests were used to compare two groups when applicable. Kruskal-Wallis tests with correction for multiple comparisons using the original FDR method of Benjamini and Hochberg were used to compare three or more groups. The exact statistical analysis methods are provided in the figure legends.

## Supporting information

Main text

## Data availability

All sequencing data have been deposited into public databases. The CITE-seq data can be found at GEO database and are available via accession number GSEXXXXXX.

## Acknowledgements

We thank all the patients and their families, as well as the nurses, pharmacists, and advanced practice practitioners who were involved in their care. SSP is generously supported by National Natural Science Foundation of China (82270157) and Zhejiang Science and Technology Program (2022C03005). CTJ is generously supported by the Nancy Carroll Allen Chair in Hematology Research, a Leukemia and Lymphoma Society SCOR grant (7020-19), and NIH R35CA242376. DAP is supported by the University of Colorado Department of Medicine Outstanding Early Career Scholar Program, the Robert H. Allen MD Chair in Hematology Research and the Leukemia and Lymphoma Society Scholar in Clinical Research Award.

## Author contributions

SSP and CTJ conceived the project and wrote the paper. SSP carried out the studies and data analysis. SSP and ITS performed xenograft experiments. SSP, BMS, and MG performed the CITE-seq experiments. AEG and KE constructed the CITE-seq dataset. AEG, SSP, and YW carried out CITE-seq data analysis. WS assisted with data visualization. SS assisted with clinical data and sample acquisitions. AI, MLA, MM, AW, SP, and TH helped with in vitro experiments. AK and TNY assisted transplant studies and in vivo treatments. JS performed morphological evaluation of bone marrow aspirates. DAP provided clinical data. DAP, CAS, and CM assisted with clinical analysis and sample acquisitions.

