## Supplementary material for "A novel type of monocytic leukemia stem cell revealed by the clinical use of venetoclax-based therapy": Main text

### Supplementary Fig. 1

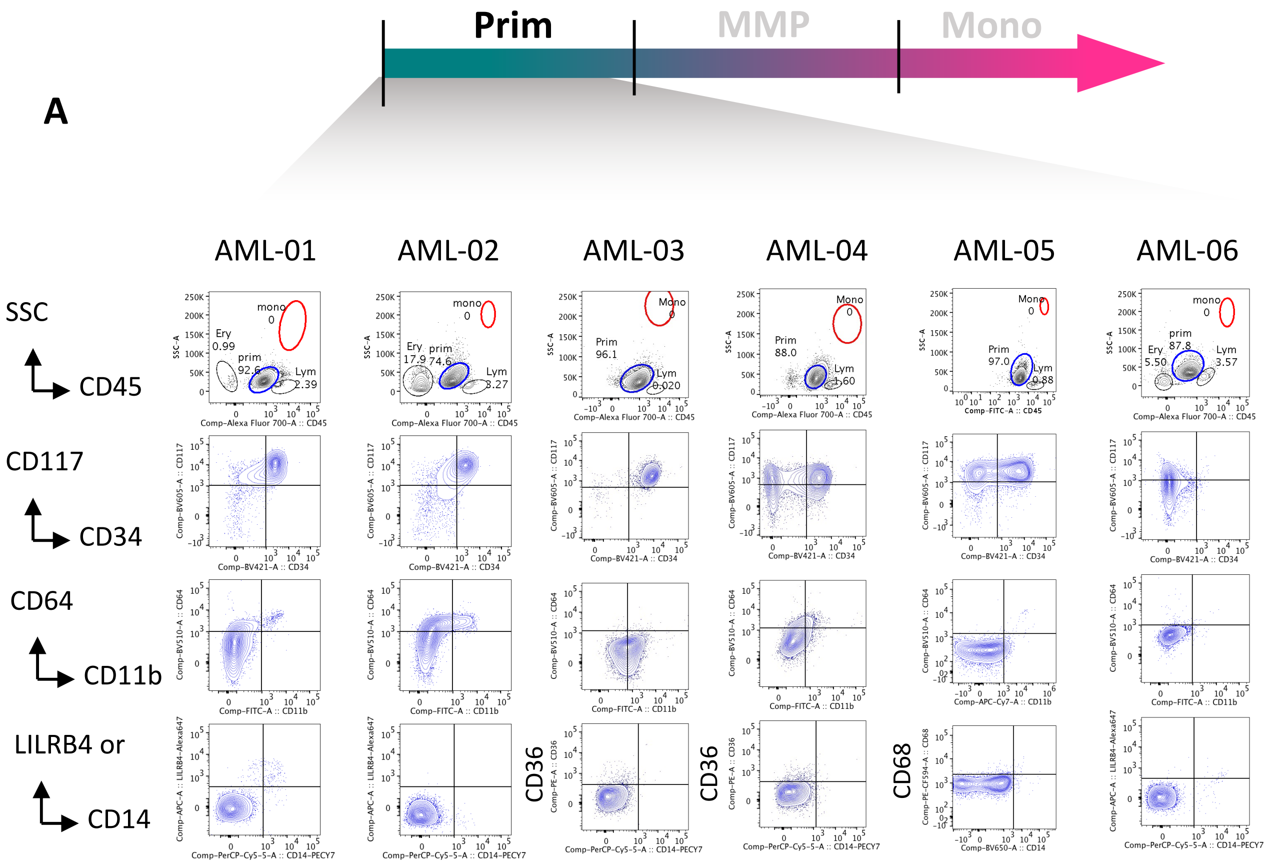

**Supplementary Fig. 1**

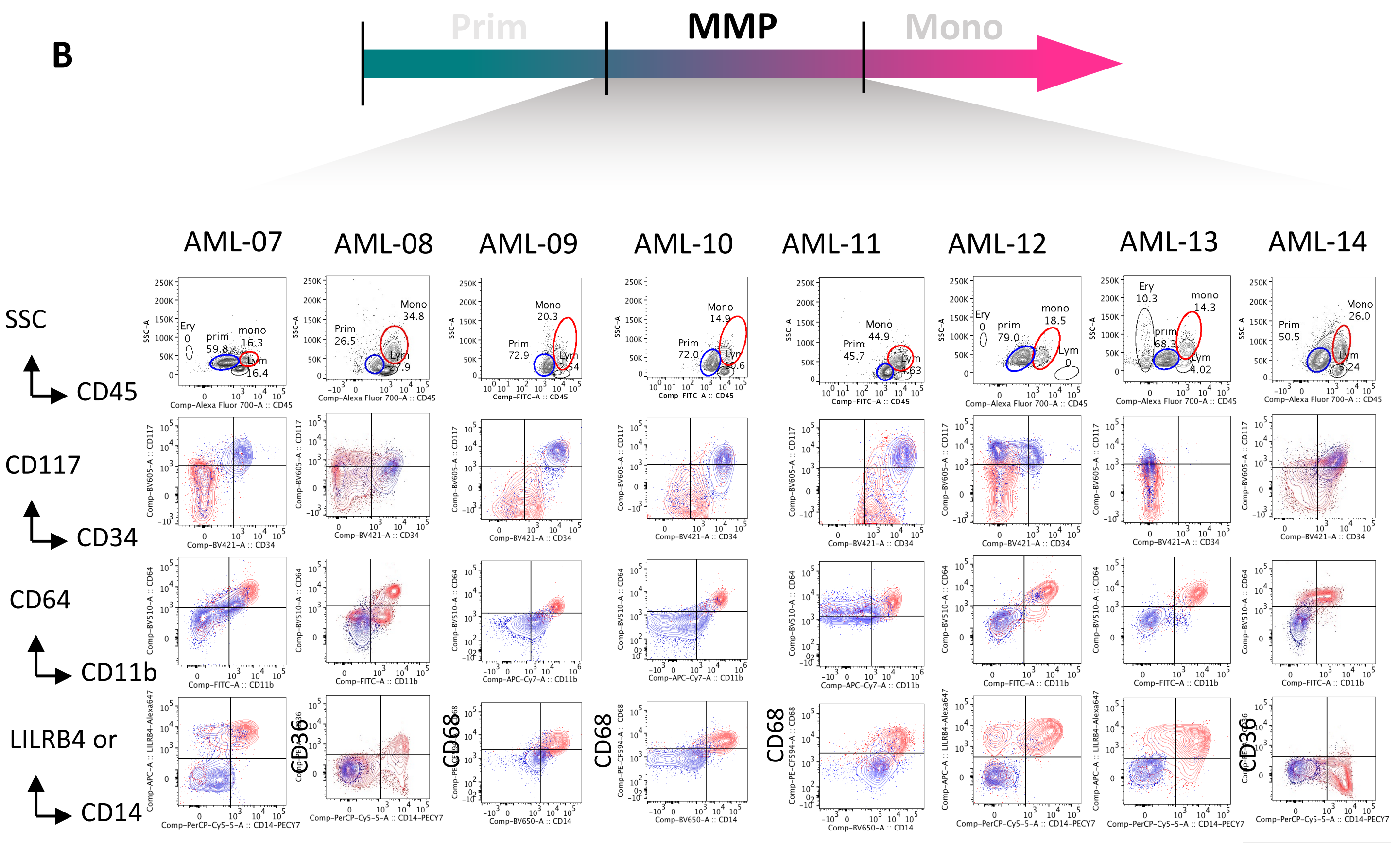

### Supplementary Fig. 1

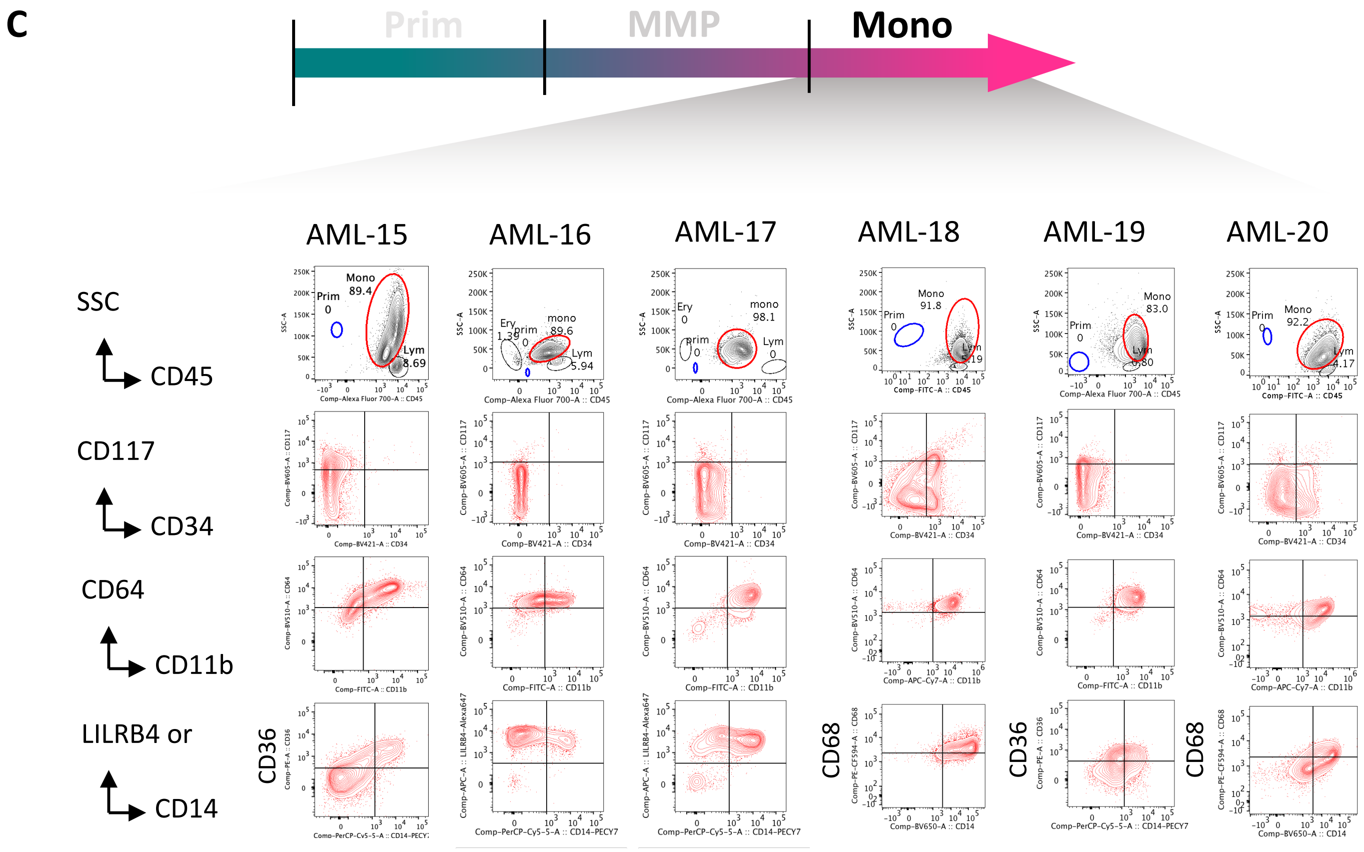

**Supplementary Fig. 1. Immunophenotyping of primary AML specimens. A-C,** Flow plots showing immunophenotyping results of Prim (**A**), MMP (**B**), and Mono (**C**) primary AML specimens. In the CD45/SSC plots, prim, mono, lym, and ery stand for primitive, monocytic, lymphocytic, and erythroid, respectively; the numbers indicate their percentages of total viable cells. In the CD34/CD117, CD11b/CD64, and CD14/LILRB4 or CD68 or CD36 plots, blue and red contours represent prim and mono cells from the CD45/SSC plots, respectively. LILRB4, CD68 and CD36 were used interchangeably to determine monocytic differentiation status

### Supplementary Fig. 2

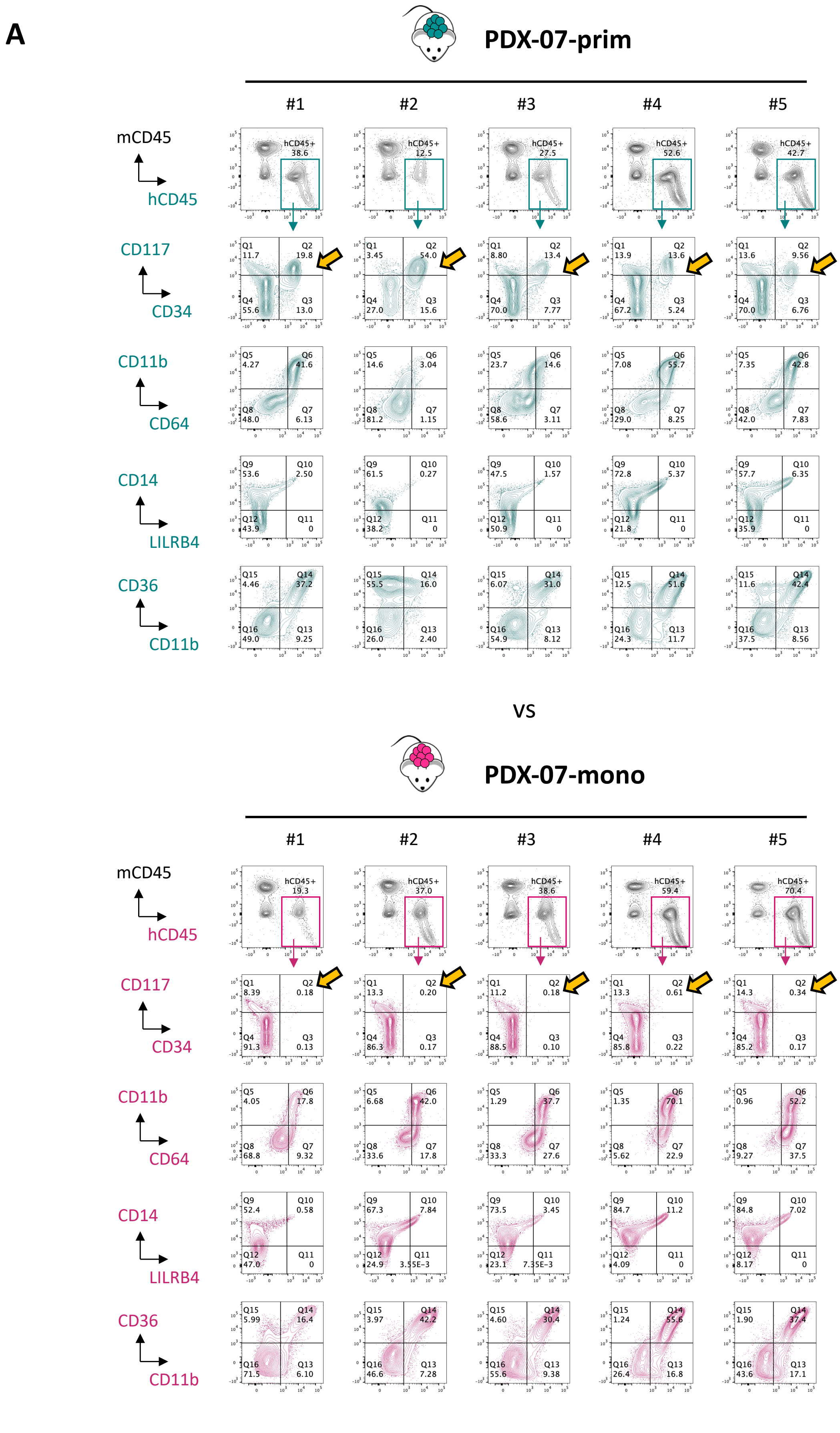

### Supplementary Fig. 2

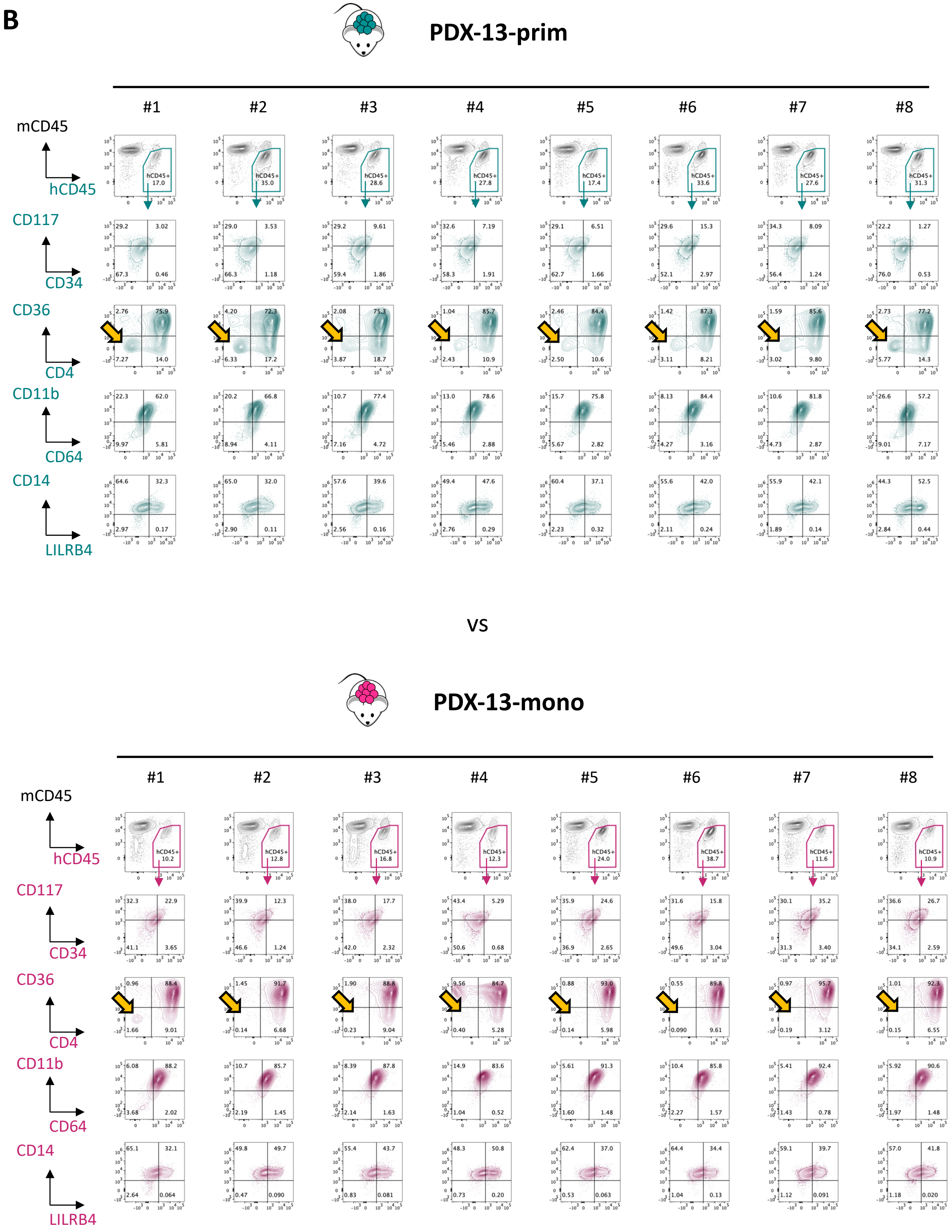

**Supplementary Fig. 2. Immunophenotyping of engrafted cells in PDX. A-B,** Flow plots showing immunophenotypic differences between leukemia arising from prim and mono subpopulations of Multi-MMP specimens AML-07 (**A**) and AML-13 (**B**) in NSG-S mice. For each mouse, engrafted human leukemic cells were gated as hCD45+/mCD45-. Subsequently, the engrafted human leukemic cells were compared with regard to their expression patterns of CD34/CD117, CD64/CD11b, LILRB4/CD14, and CD11b/CD36. Teal and pink contour plots represent prim- and mono-engrafted cells, respectively. Yellow arrows highlight the most distinct differences between the groups.

### Supplementary Fig. 3

**A**

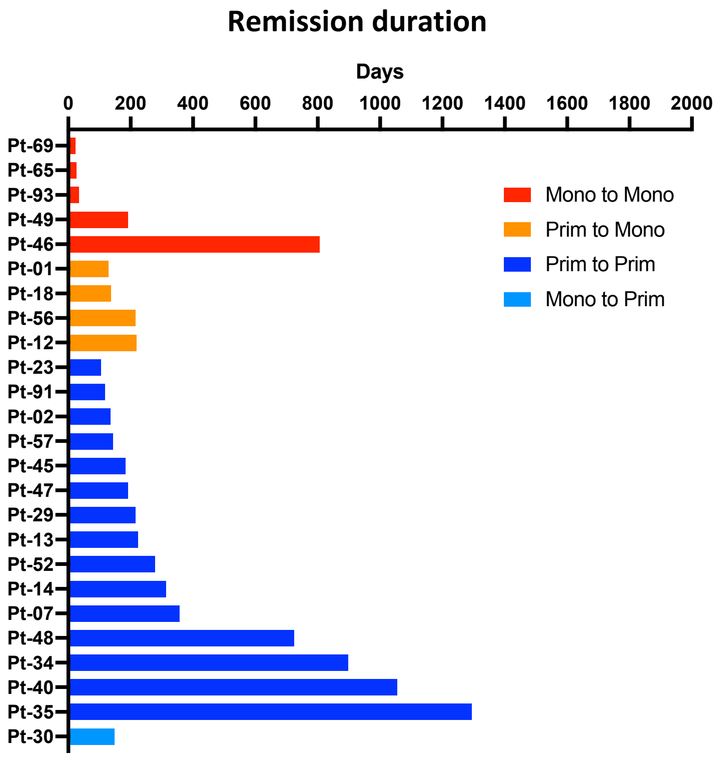

**Supplementary Fig. 3. Remission duration of VEN/AZA relapsed AML patients. A,** Bar graphs showing remission duration in days for a cohort of 25 AML patients who received the VEN/AZA therapy and experienced relapse response. The cohort contains five patients who sustained a monocytic phenotype between diagnosis and relapse (red bars, mono to mono), four patients transited from a primitive phenotype at diagnosis to monocytic phenotype at relapse (orange bars, prim to mono), 15 patients sustained a primitive phenotype between diagnosis and relapse (blue bars, prim to prim), and one patient transited from a monocytic phenotype at diagnosis to primitive phenotype at relapse (light blue bar, mono to prim).

### Supplementary Fig. 4

## A

Lineage-specific surface antigens (protein/adt)

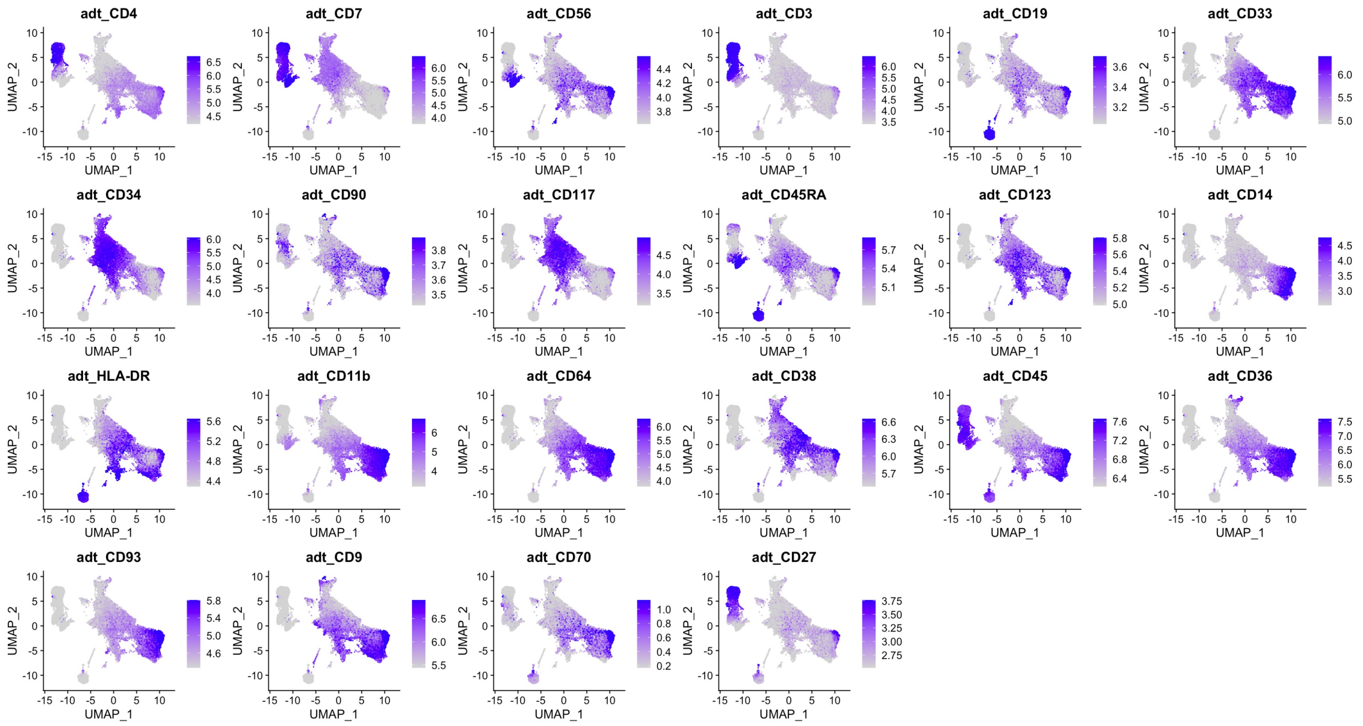


## B

Lineage-specific surface antigens (RNA)

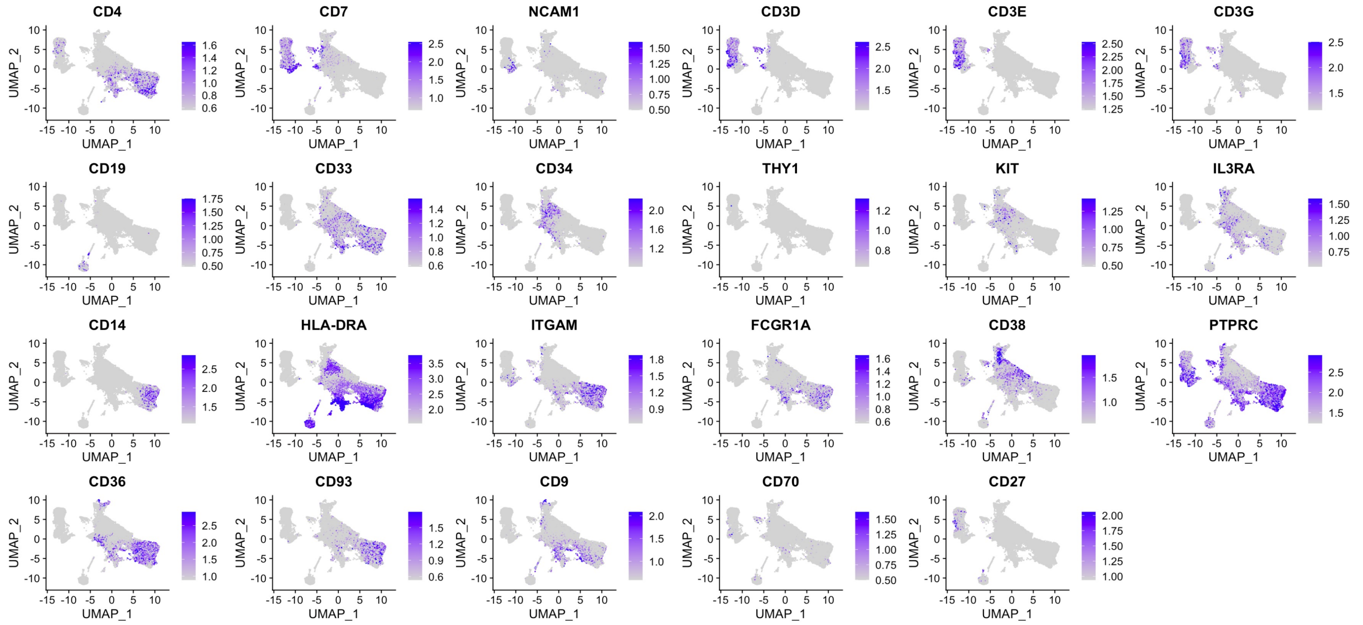

### Supplementary Fig. 4

**C**

Lineage specific transcription factors (RNA)

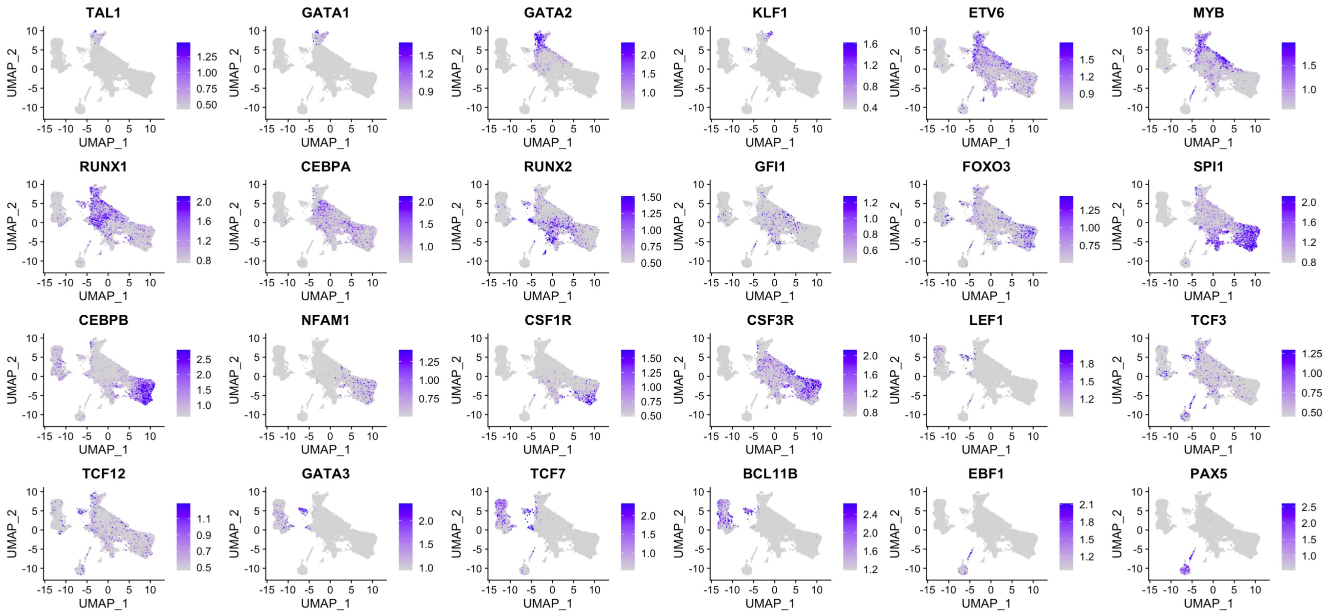


**D**

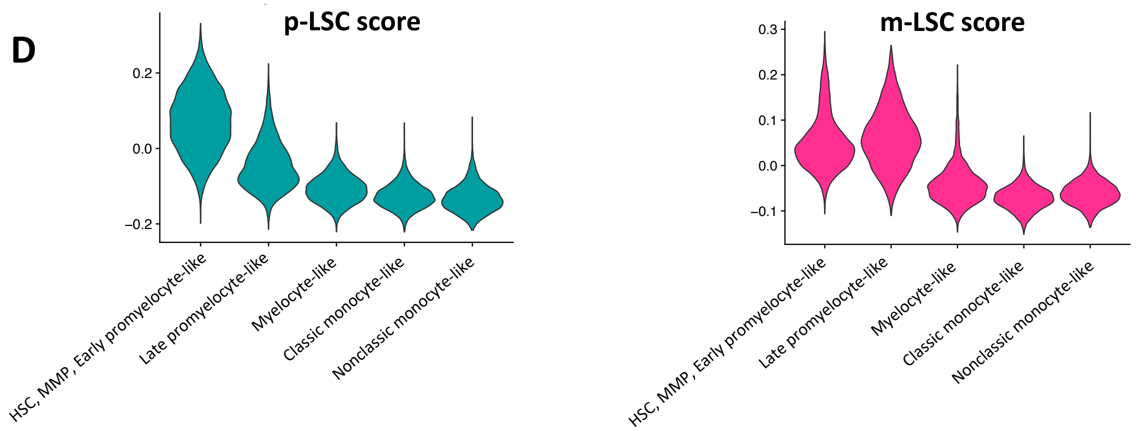


**E**

**Up-regulated in p-LSC**

**Up-regulated in m-LSC**

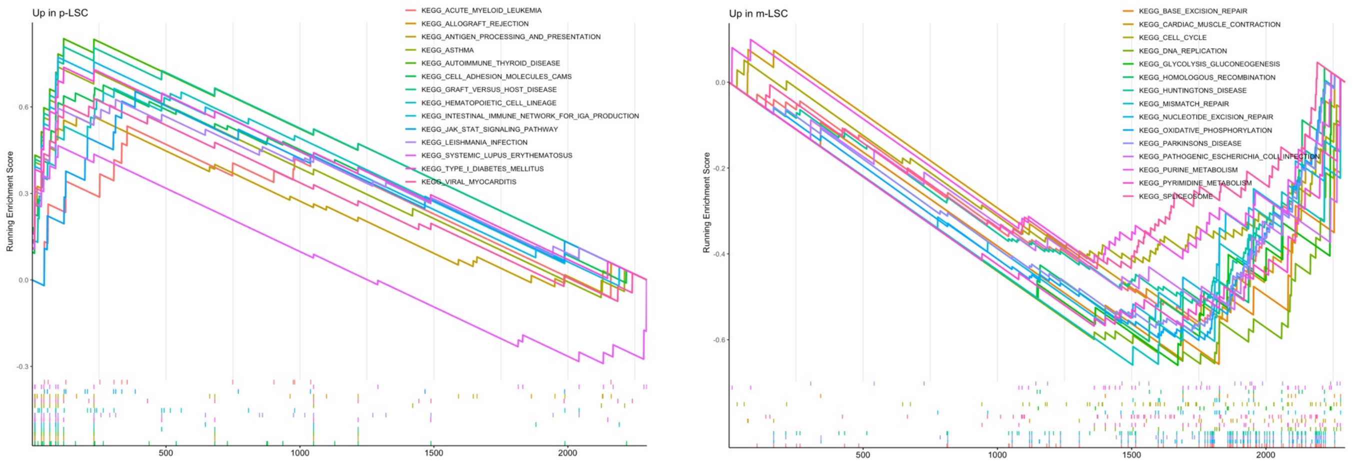

**Supplementary Fig. 4. CITE-seq data analysis. a-b,** Protein (**A**) and RNA (**B**) expression of lineage-specific surface antigens on UMAP. **C,** RNA expression of lineage-specific transcription factors on UMAP. **D,** Violin plots showing distribution of p-LSC and m-LSC scores on cells of HSPC, MPP, Early Promyelocyte-like, Late promyelocyte-like, Myelocyte-like, classical monocyte-like, and Nonclassical monocyte-like subclusters. **E,** GSEA enrichment plots showing top upregulated KEGG pathways in p-LSC and m-LSCs.

### Supplementary Fig. 5

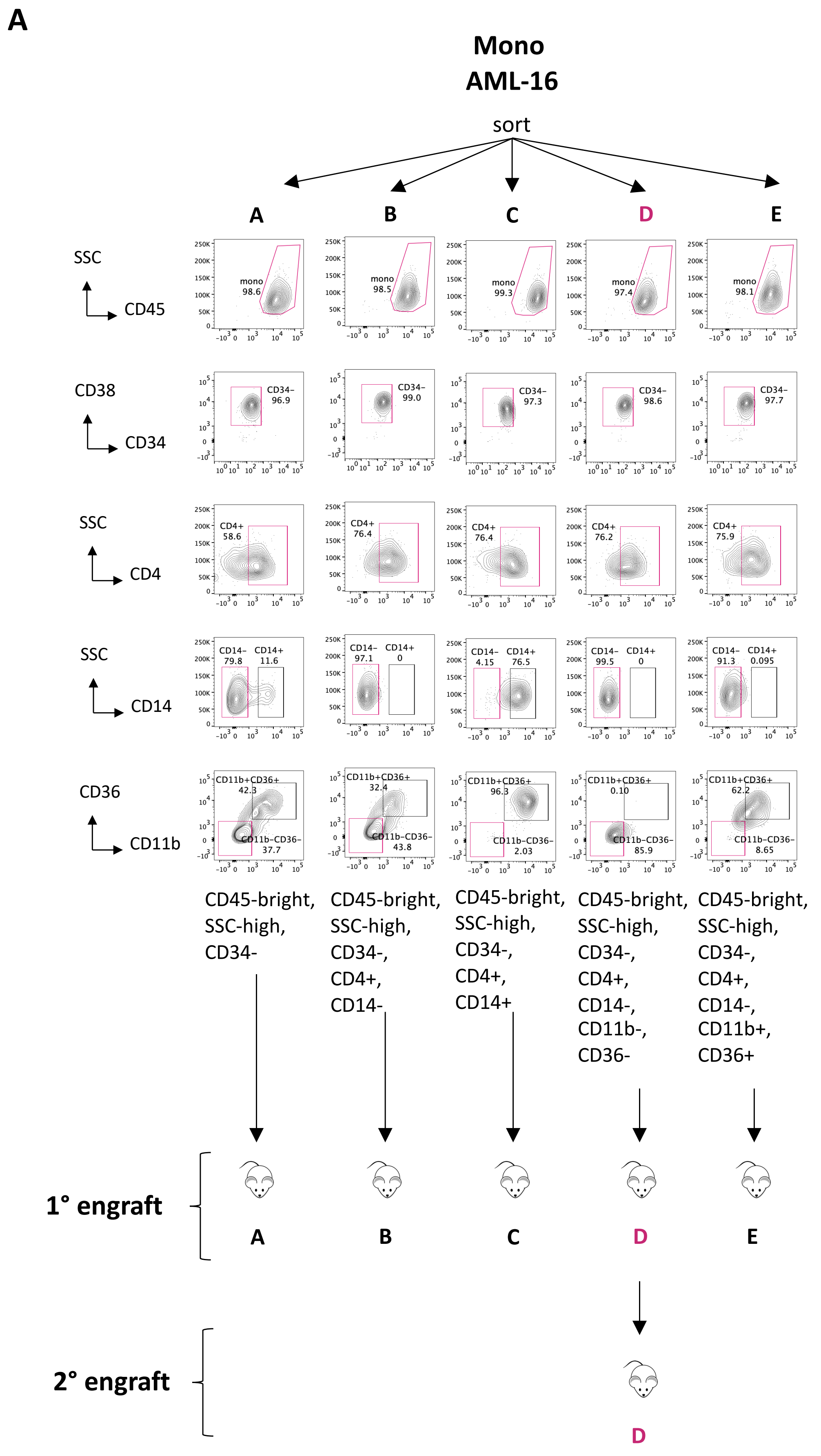

### Supplementary Fig. 5

**B**

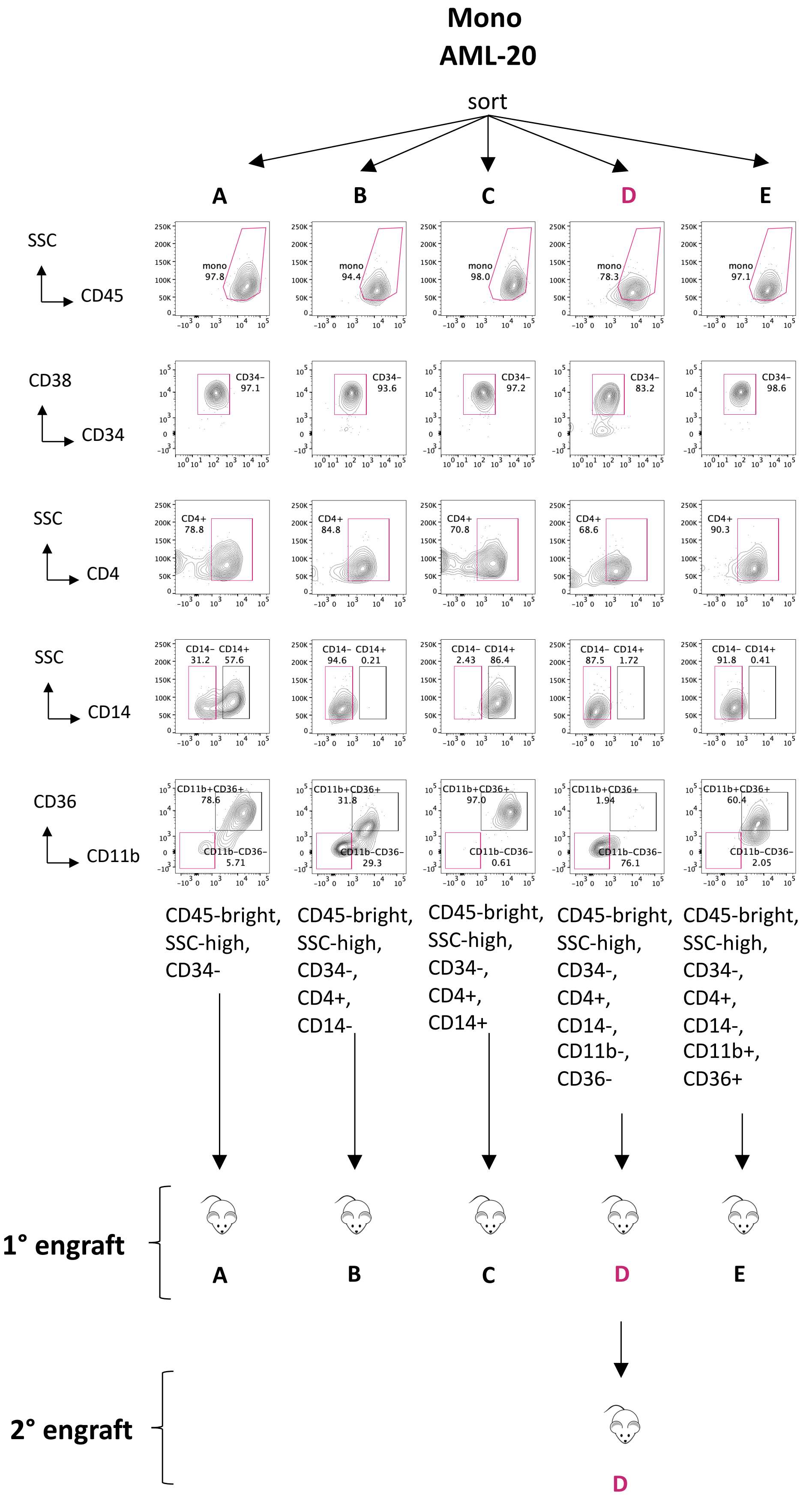

### Supplementary Fig. 5

C

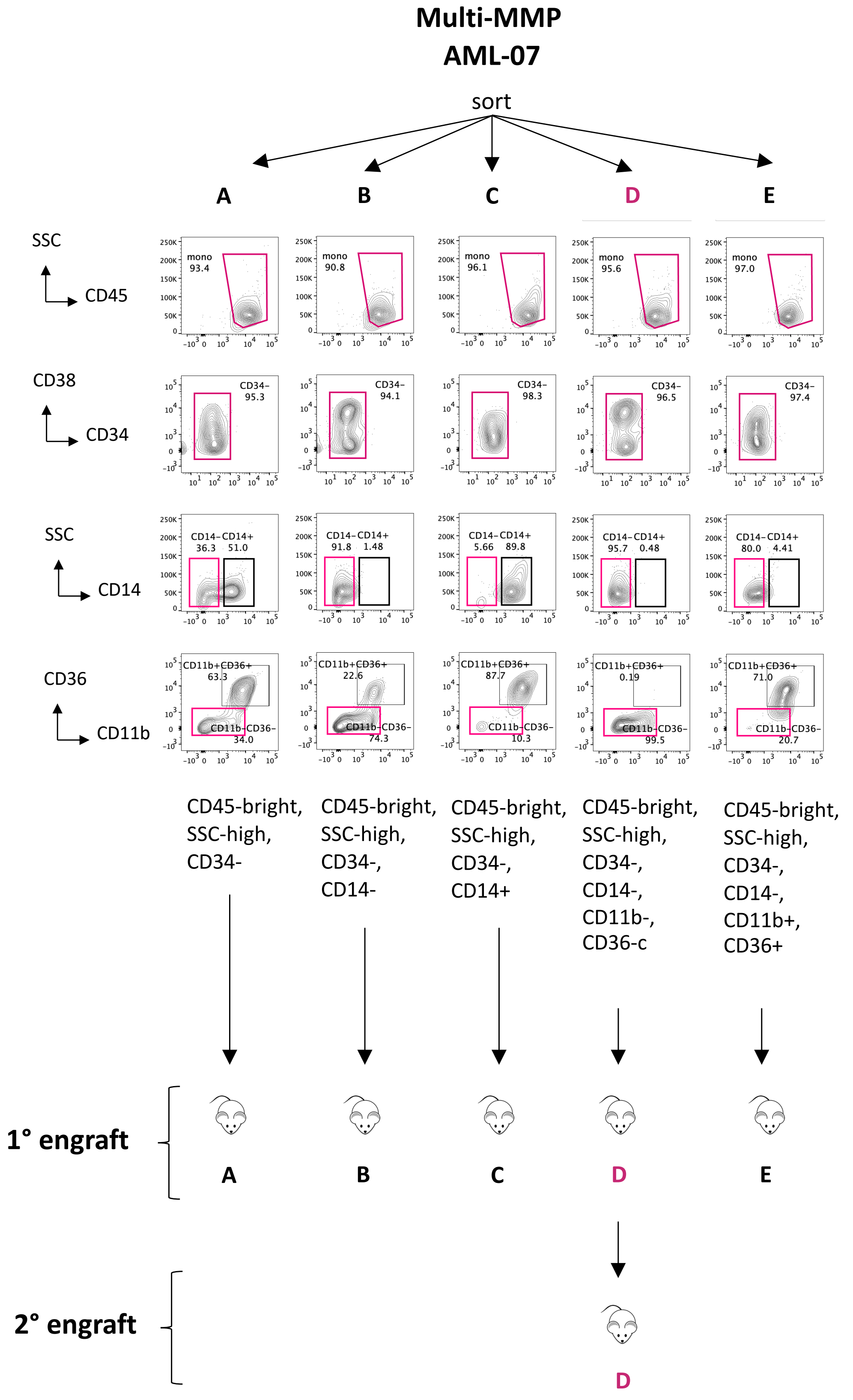

### Supplementary Fig. 5

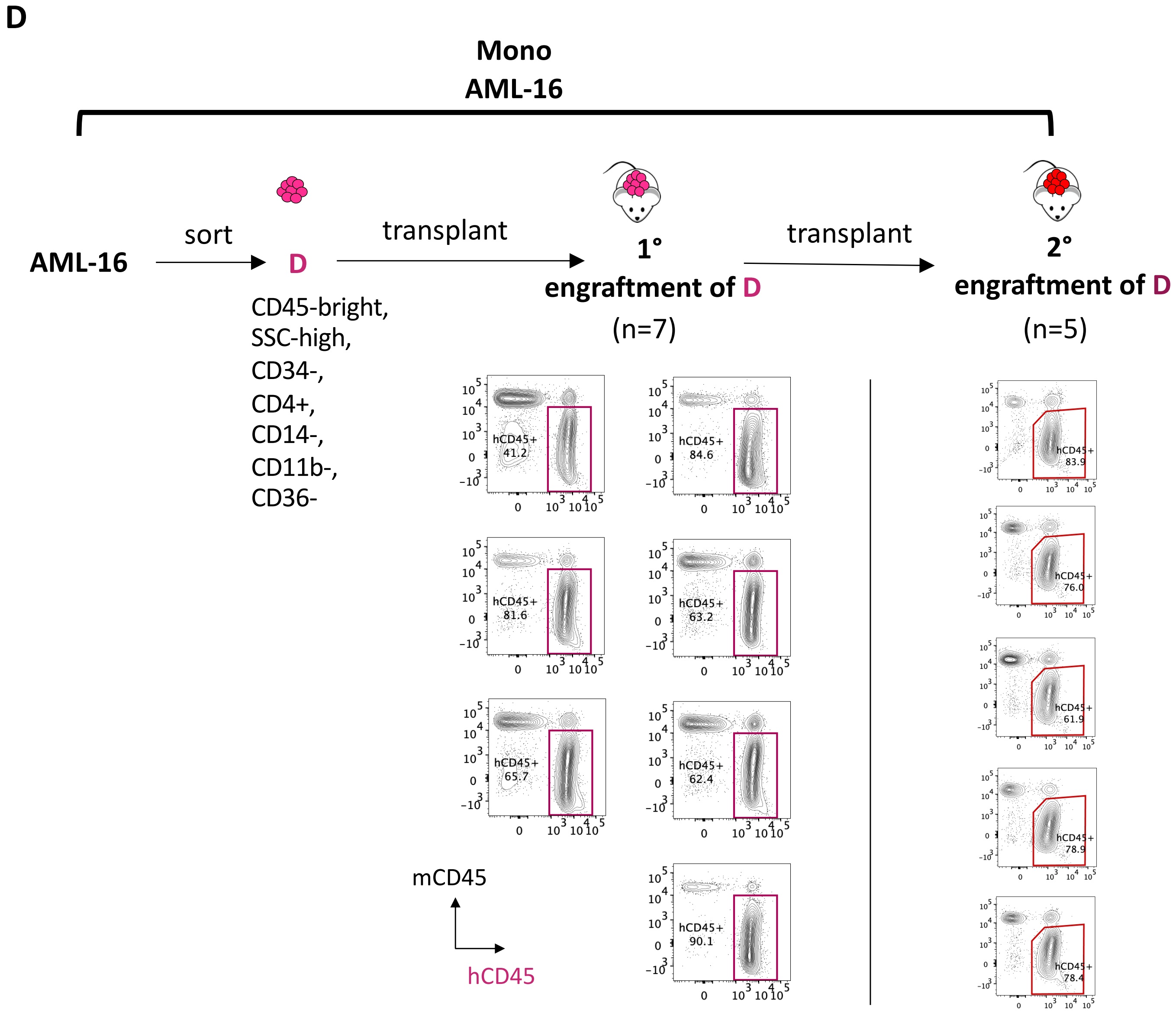

#### Supplementary Fig. 5

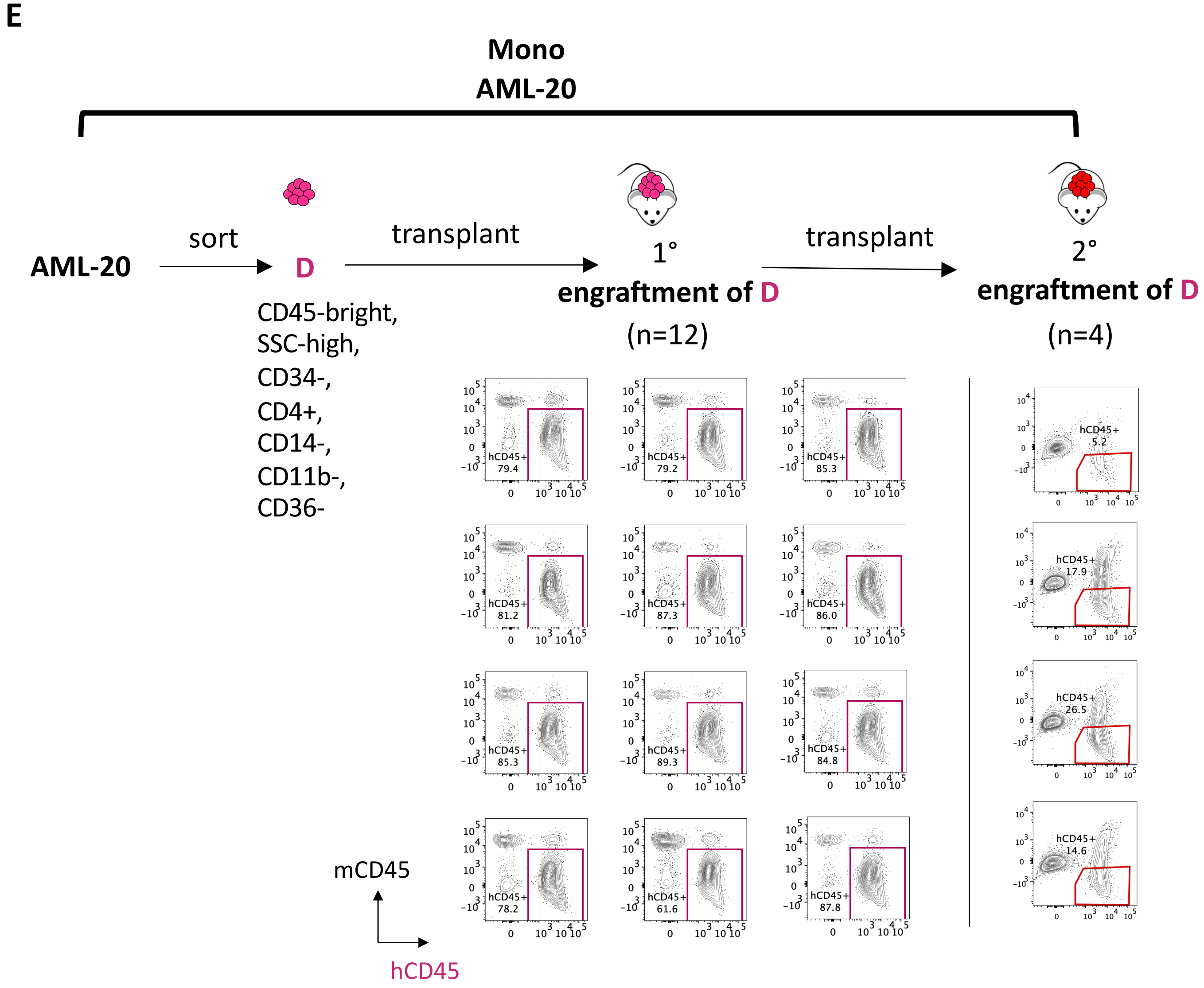

### Supplementary Fig. 5

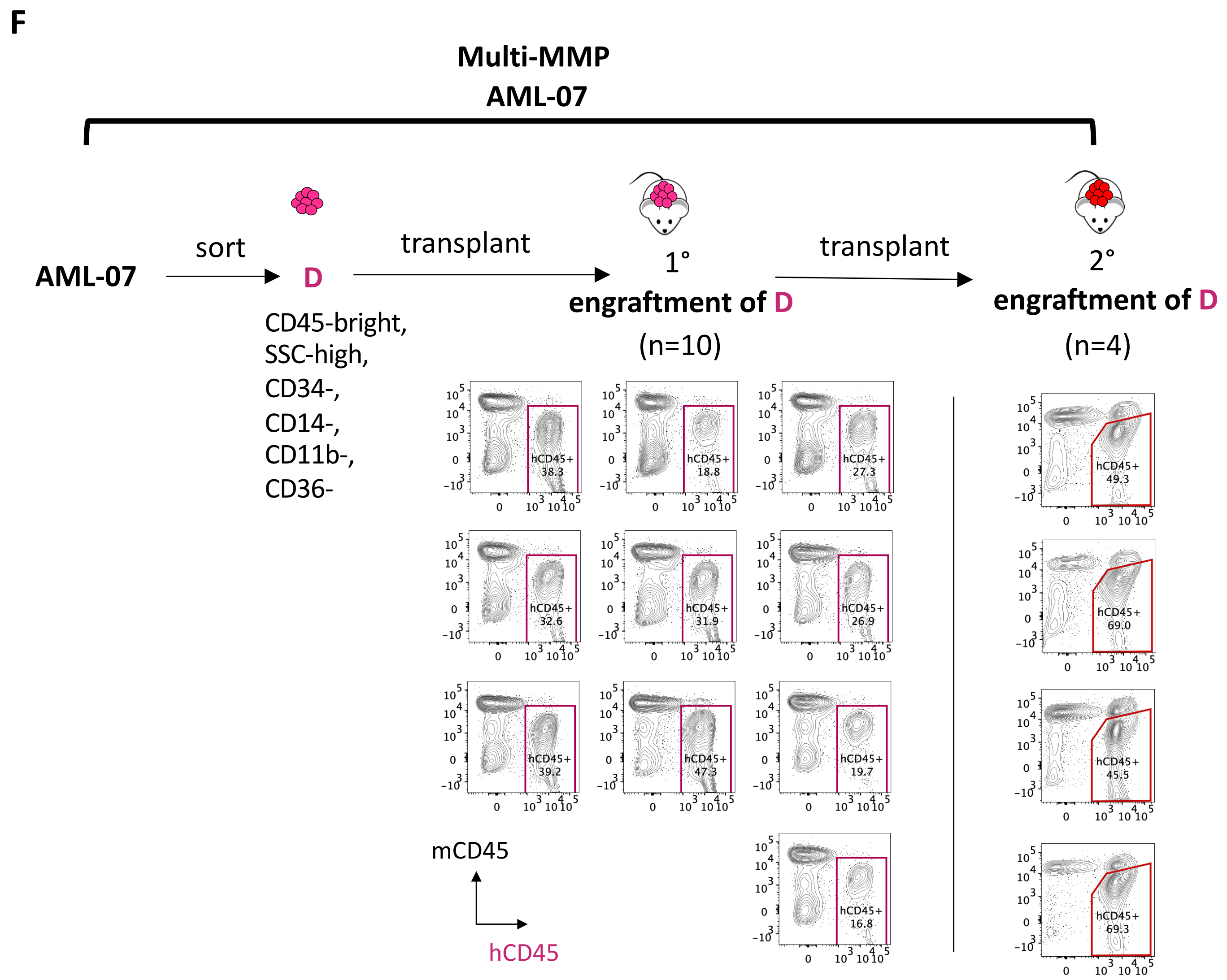

### Supplementary Fig. 5

## G

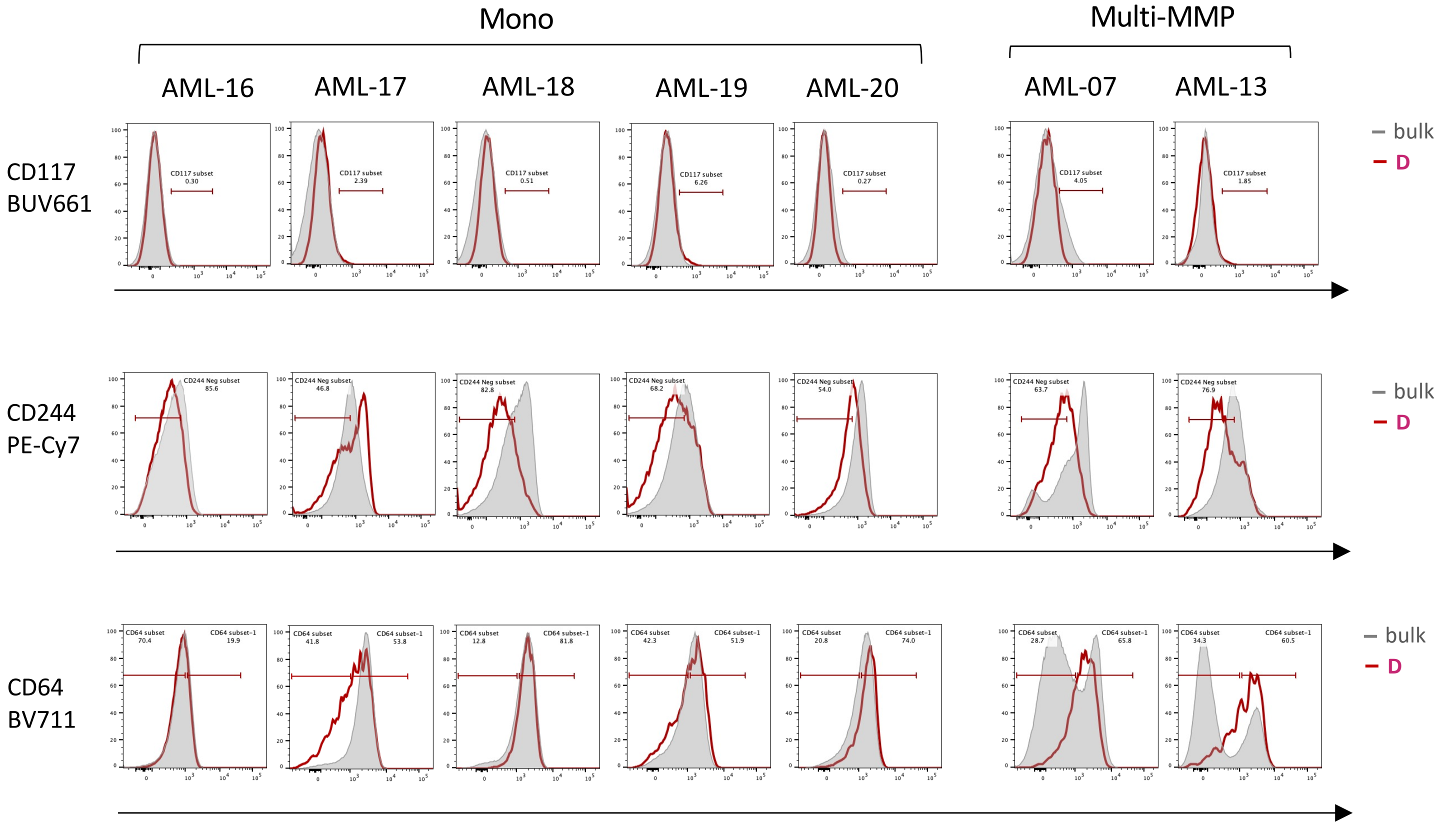

**Supplementary Fig. 5. Sorting strategies for determining m-LSC immunophenotype. A-C,** Immunophenotyping of A, B, C, D, and E subpopulations sorted from Mono AML-16 (**A**), Mono AML-20 (**B**), and Multi-MMP AML-07 (**C**), and used for injection into NSG-S mice to determine their m-LSC potential. Red gates highlight cells with enriched m-LSC potential, Black gates highlight the cells without m-LSC activity. Texts in the bottom of flow plots summarize the exact immunophenotype of each group injected into NSG-S mice. **D-F,** Tumor burden in primary and secondary transplants shown by human CD45 and mouse CD45 staining. **G,** Expression of CD117, CD244, and CD64 in m-LSC enriched D vs bulk populations.

**Supplementary Fig. 6**

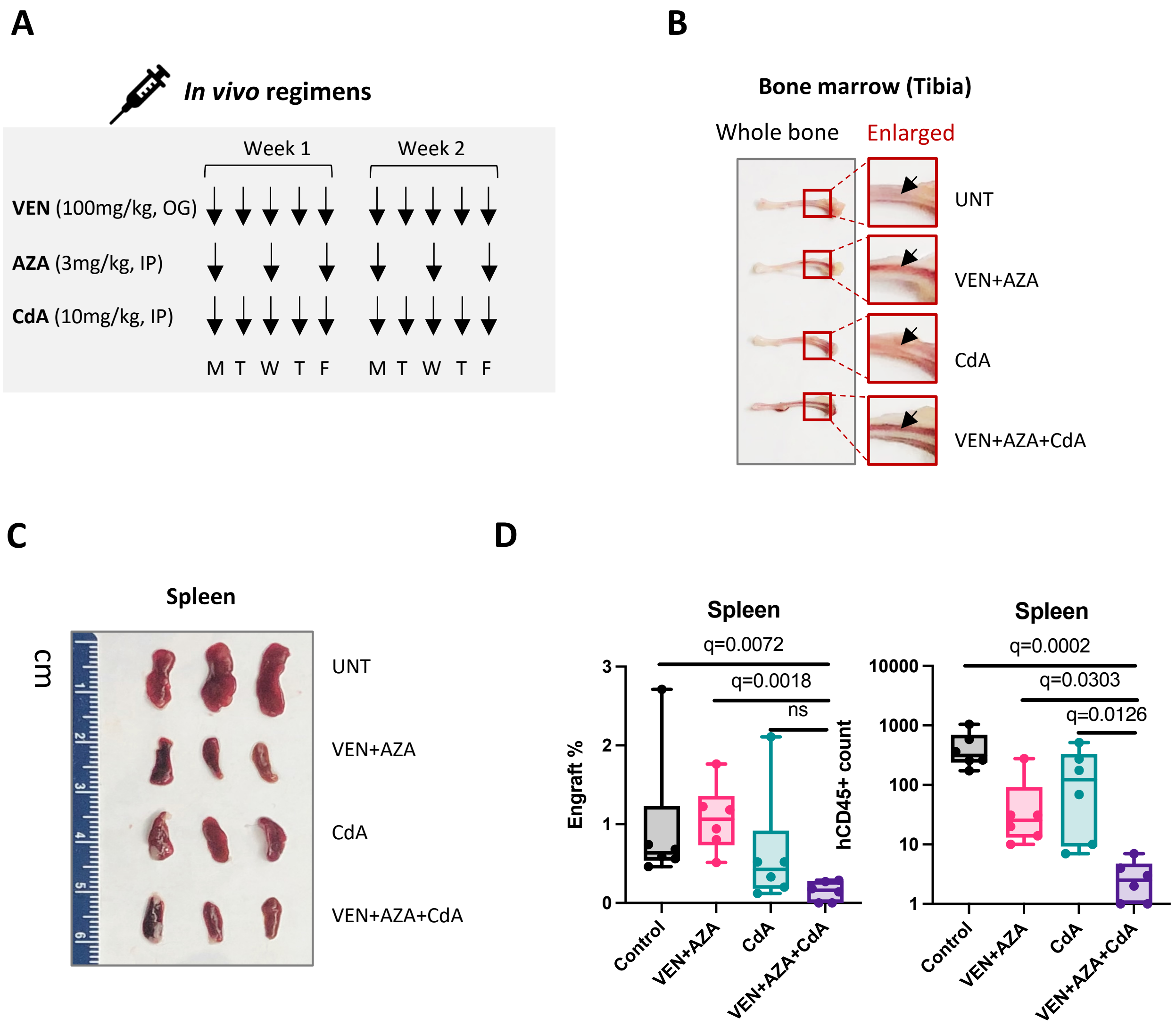

**Supplementary Fig. 6. In vivo efficacy of cladribine (CdA) in combination with VEN+AZA in treating a PDX model of Multi-MMP AML. A,** Design of VEN, AZA, CdA regimens. **B,** Images of representative tibia bones from all treatment groups showing clearance of leukemia burden as the leukemia-filled pale white marrow space become restored as the healthier red marrow space in the VEN+AZA and even more in the VEN+AZA+CdA group. **C,** Images of representative spleens from all treatment groups showing restoration of normal spleen size in treated groups. **D,** Impact of VEN+AZA, CdA or combo treatments on the spleen tumor burden of PDX. Engraft% was determined by % of hCD45+/mCD45- cells within total viable spleen mononuclear cells. hCD45+ count was determined by direct quantification of hCD45+/mCD45- cells within a set volume of spleen harvest using flowcytometry. Each dot represents a unique mouse. Control (n=6), VEN+AZA (n=6), CdA (n=6), VEN+AZA+CdA (n=6). Box plots represent median +/- interquartile. Kruskal-Wallis test was used. ns, not significant.

### Supplementary Fig. 7

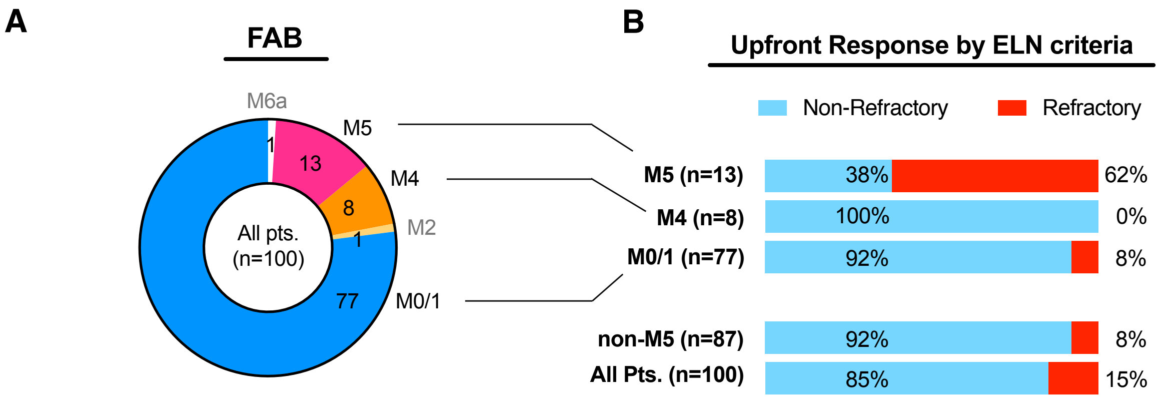

**Supplementary Fig. 7. M5 but not M4 patients showing significantly higher refractory rate to VEN/AZA therapy. A,** A circular pie chart showing numbers of patients identified in different FAB subclasses (Reproduction of Pei et al, Cancer Discovery, 2020 Supplementary Figure S1A). **B,** Bar graphs showing percentage of patients had refractory or non-refractory responses to VEN+AZA therapy according to the ELN criteria. (Reproduction of Pei et al, Cancer Discovery, 2020 Supplementary Figure S1B).

#### Supplementary Tables

**Supplementary Table S1.** Clinical information of primary AML specimens

**Supplementary Table S2.** Clinical information of specimens from VEN/AZA relapsed AML patients

**Supplementary Table S3.** CITE-seq and flow antibodies

**Supplementary Table S4.** CITE-seq specimens

**Supplementary Table S5.** p-LSC and m-LSC gene expression signatures

**Supplementary Table S6.** Upregulated genes in p-LSCs

**Supplementary Table S7.** Upregulated genes in m-LSCs

**Supplementary Table S8.** Results of GSEA analysis comparing p-LSCs to m-LSCs

**Supplementary Table S9.** PDX experiments
